# Valine Engages a Prefrontal mGluR5 Allosteric Site to Preserve Social Behavior

**DOI:** 10.64898/2026.10.14.759089

**Authors:** Xingyue Shan, Tailin Zhu, Yuwei Wang, Tian-Tian Shi, Xiaoyu Yang, Susu Wang, Xin Zhang, Wei Zhou, Zeyu Zhou, Haoran Geng, Jiangyu Yan, Hui Lv, Yuhua Yan, Yue Guo, Lan Shi, Lingli Zhang, Keyi Hu, Dandan Wang, Ye Yu, Bing Mei, Qianlong Zhang, Haipeng Sun, Xiaoqun Wang, Qianhui Qu, Jian-Yuan Zhao, Wei-Guang Li, Fei Li

**Author notes:** Correspondance (X.W.), (Q.Q.), (J.-Y.Z.), (W.-G.L.), (F.L.). These authors contributed equally (Xingyue Shan, Tailin Zhu, Yuwei Wang, Tian-Tian Shi, Xiaoyu Yang, Susu Wang).

## Abstract

Branched-chain amino acid metabolism is linked to neurodevelopmental disorders, but whether a defined amino acid directly modulates cortical receptor signaling to control behavior is unknown. Here, we show that valine is uniquely required for sociability through medial prefrontal cortex (mPFC) circuitry. Partial dietary valine restriction, but not comparable leucine or isoleucine restriction, or mPFC-restricted acceleration of BCAA catabolism through Bckdk loss, disrupts social interaction, and local valine supplementation rescues behavior. LiP-SMap, surface plasmon resonance, cryo-EM, and mutagenesis identify metabotropic glutamate receptor 5 (mGluR5) as a direct valine target and reveal a valine-selective allosteric pocket distinct from the glutamate site. Valine is signaling-biased: it drives β-arrestin-linked output yet has little standalone Gq efficacy, instead potentiating glutamate-evoked mGluR5 signaling to recruit endocannabinoid/CB1 feedback, gate inhibitory transmission, and preserve mPFC-dependent social behavior. Together, these findings identify valine as a diet-derived allosteric ligand that links amino acid availability to cortical circuit control of social behavior.

**Graphical abstract:** 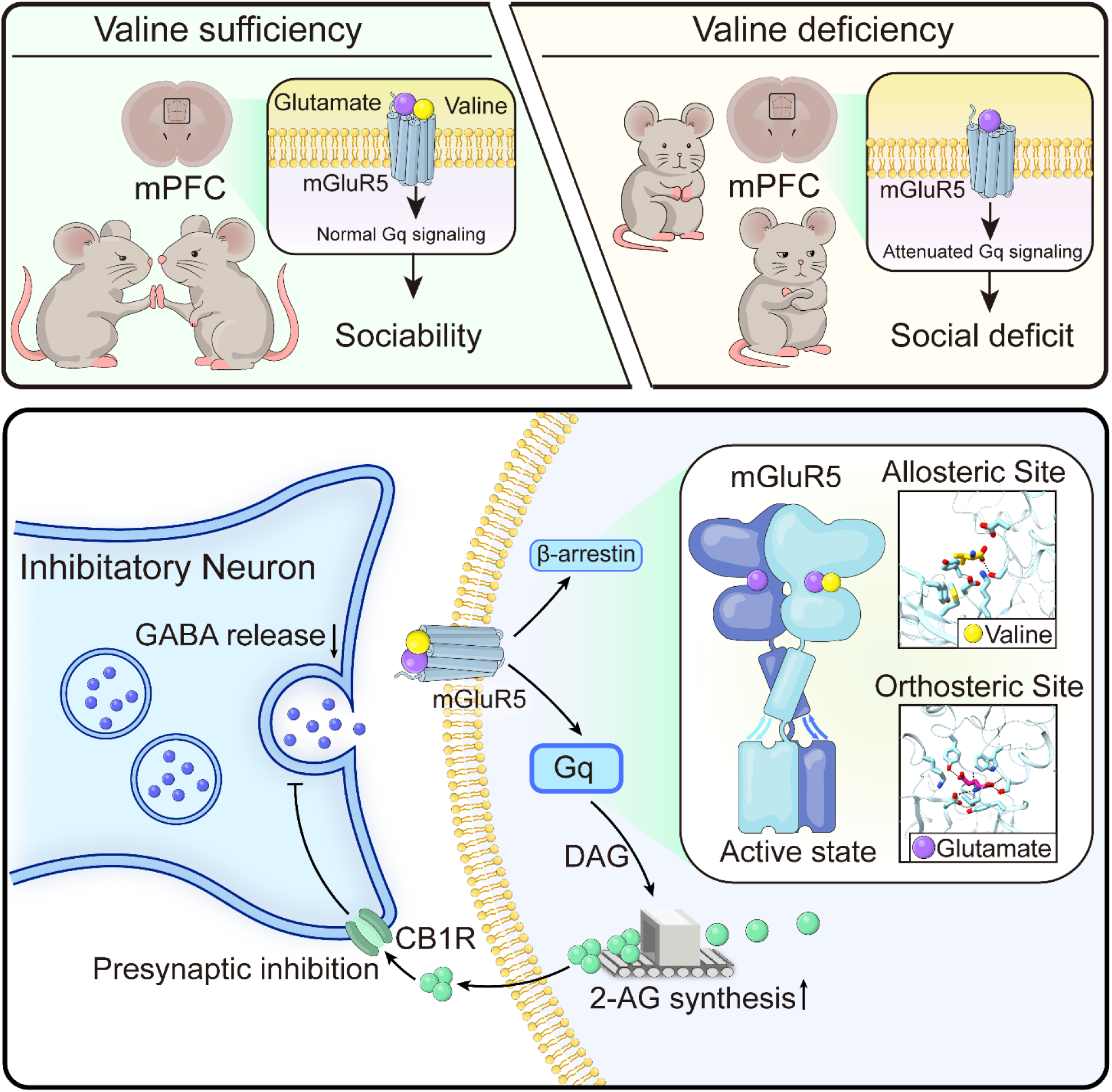

**In Brief:** Valine, an essential amino acid, directly engages an allosteric pocket in mGluR5 and biases receptor signaling in the medial prefrontal cortex. By potentiating glutamate-driven mGluR5–endocannabinoid suppression of inhibition, valine preserves excitation–inhibition balance and social behavior.

**Highlights:**

1. Valine is uniquely required in mPFC to support sociability.
2. LiP-SMap, SPR, and cryo-EM identify a valine pocket in mGluR5.
3. Valine biases mGluR5 signaling and potentiates glutamate-evoked output.
4. Valine–mGluR5 recruits CB1 feedback to gate inhibition during behavior.

## Introduction

Branched-chain amino acids (BCAAs)—valine, leucine, and isoleucine—are essential nutrients whose brain levels reflect a dynamic interplay between diet, blood–brain barrier (BBB) transport, and regulated catabolic flux.^1–6^ In neural tissue, BCAAs support biosynthetic and energetic demands, contribute nitrogen to neurotransmitter-related metabolism, and compete for entry into the brain through large neutral amino-acid transport systems at the BBB.^7–10^ Consequently, even modest perturbations in BCAA availability can alter the biochemical state of the brain. A central control point in this architecture is the branched-chain α-ketoacid dehydrogenase (BCKDH) step, which is restrained by branched-chain ketoacid dehydrogenase kinase (BCKDK), positioning BCKDK as a key regulator of local BCAA tone.^11,12^ These features make BCAAs plausible determinants of brain function not only through long-term effects on nutrition and growth, but also through local, region- and cell-type–specific metabolic control.

Consistent with this possibility, perturbed amino acid profiles—including BCAA-related signatures—have been associated with neurodevelopmental phenotypes. Metabolomics studies have identified autism spectrum disorder (ASD) subgroups enriched for amino-acid dysregulation, although cohort heterogeneity has made it difficult to infer mechanism from association alone.^13,14^ Human genetics provides a stronger causal foothold: loss-of-function mutations in *BCKDK* have been identified in families with ASD, epilepsy, and intellectual disability, implicating dysregulated BCAA catabolism as a driver of neurodevelopmental dysfunction.^15,16^ In parallel, disruption of BBB transport of large neutral amino acids through SLC7A5 reduces brain BCAA availability and produces neurological and behavioral abnormalities accompanied by synaptic changes consistent with amino-acid deprivation.^17^ Notably, BCAA-linked phenotypes extended beyond social behavior to include motor and developmental deficits,^18^ arguing against a single “global nutrition” explanation and instead suggesting that BCAA availability may be decoded in a circuit- and region-specific manner. Yet mechanistic interpretations in the field remain dominated by indirect frameworks—nutrient sensing and proteostasis pathways such as mTORC1 and GCN2, altered protein synthesis, or broad developmental compromise—that can explain pervasive phenotypes but rarely identify direct molecular effectors with synaptic specificity.^19^ At the same time, a broader conceptual possibility is emerging: small metabolites can directly engage proteins to alter signaling states, and G-protein coupled receptors (GPCRs) can be modulated by endogenous ligands beyond canonical neurotransmitters. Whether an essential amino acid is directly decoded at cortical synapses by a defined receptor-level mechanism, and whether such a mechanism can shape behavior on circuit-relevant timescales, remains largely unknown.

We addressed this problem in the context of social interaction, a behavioral domain that depends strongly on medial prefrontal cortex (mPFC) function and on tight regulation of synaptic gain and excitation–inhibition balance.^20–25^ We asked three related questions. Are BCAAs functionally interchangeable in supporting sociability, or does one of them act as a behaviorally instructive signal within mPFC? If such specificity exists, can it be explained by an acute receptor-level decoding mechanism rather than by broad metabolic deprivation? And if so, how does this mechanism reshape synaptic transmission and circuit recruitment during social behavior?

Here, we show that these questions converge on a single pathway. Valine, but not leucine or isoleucine, is uniquely required for mPFC-dependent sociability: partial dietary valine restriction impairs social interaction, mPFC-restricted acceleration of BCAA catabolism through Bckdk loss phenocopies this deficit, and acute valine supplementation restores behavior. Mechanistically, we identify metabotropic glutamate receptor 5 (mGluR5)—a key regulator of synaptic transmission and plasticity^26,27^—as a direct molecular effector through which valine is decoded in prefrontal circuits. Using chemoproteomics, biophysical binding, cryo-EM, and functional validation, we find that valine engages a selective allosteric pocket on mGluR5, biases receptor output, and potentiates glutamate-evoked signaling. In mPFC, this valine–mGluR5 coupling recruits endocannabinoid/CB1 feedback to gate inhibitory circuitry during social interaction, thereby establishing a direct molecular route by which amino acid availability can shape synaptic computation and preserve social behavior.

## Results

### Local disruption of BCAA catabolic control in mPFC identifies valine as uniquely required for sociability

To determine whether a specific BCAA supports social behavior, and whether such a requirement can be localized to mPFC, we combined systemic dietary perturbations with a region-restricted strategy to accelerate BCAA catabolism in mPFC. We first fed mice diets in which a single BCAA—valine, leucine, or isoleucine—was reduced by 50% for 4 weeks (Figure 1A). Cerebrospinal fluid (CSF) metabolomics separated the diet groups by principal component analysis (PCA) (Figure 1B), and targeted quantification confirmed that each diet selectively lowered the corresponding BCAA in CSF (Figures 1C–1E). In the three-chamber sociability assay, control mice robustly preferred the chamber containing a social partner (S1) over the empty chamber (EM). Strikingly, 50% valine restriction abolished sociability, eliminating the normal preference for S1, whereas comparable restriction of leucine or isoleucine preserved chamber preference (Figures 1F–1H). Thus, within the BCAA class, sociability shows a specific dependence on valine rather than a generic sensitivity to partial BCAA restriction.

**Figure 1.**
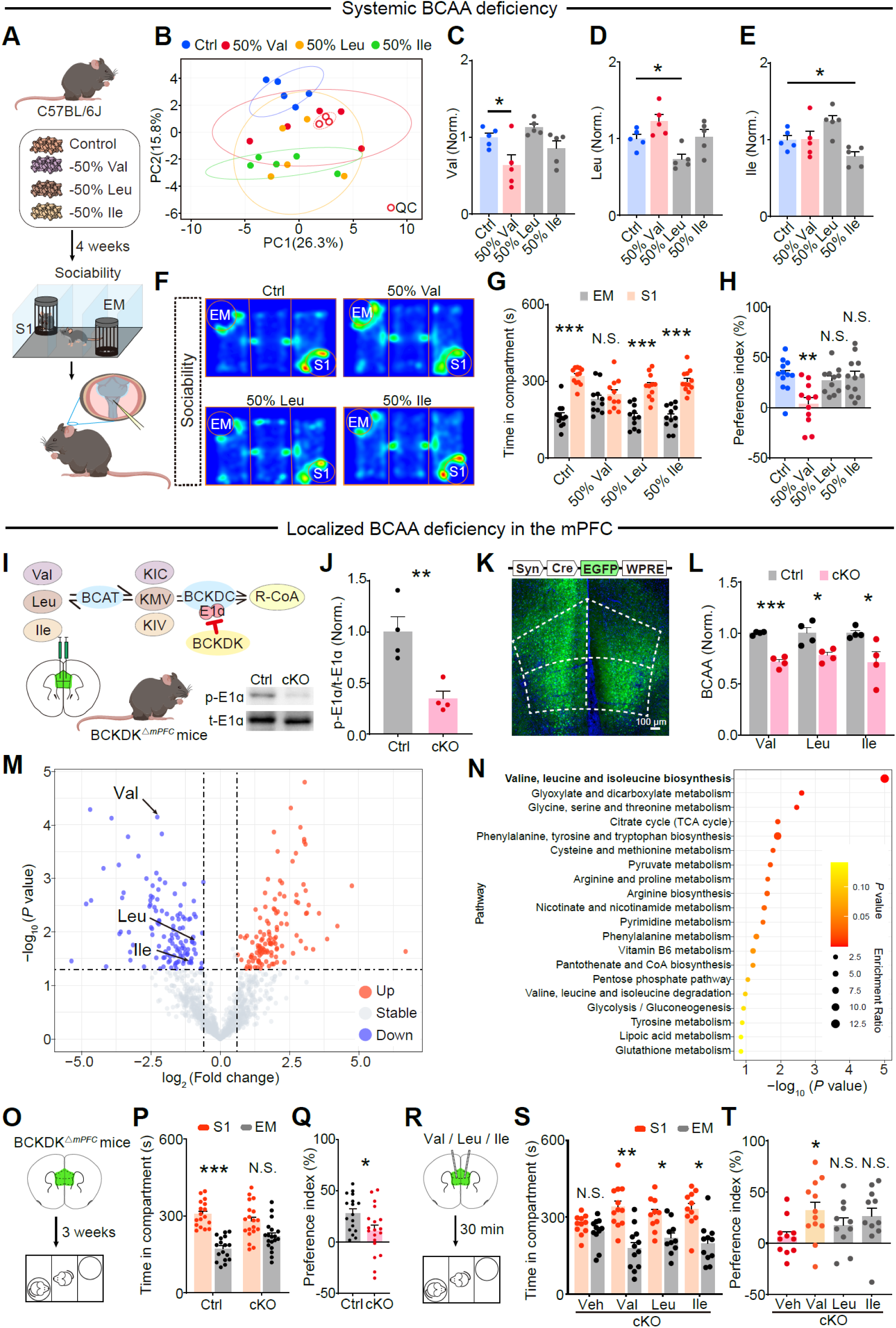
Valine is selectively required in mPFC for sociability. (A) Experimental design for systemic BCAA manipulation. C57BL/6J mice were maintained on control chow or diets in which a single BCAA [valine (Val), leucine (Leu), or isoleucine (Ile)] was reduced by 50% for 4 weeks, followed by three-chamber sociability testing and cerebrospinal fluid (CSF) collection for metabolomics. (B) Principal component analysis of CSF metabolomics across diet groups, with pooled quality-control (QC) samples indicated. *n* = 5 biologically independent mice per group. (c–e) CSF abundance of valine (C), leucine (D) and isoleucine (E) after dietary manipulation, normalized to control. *n* = 5 mice per group. (F) Representative occupancy heat maps during the sociability phase. EM, empty chamber; S1, stranger 1. (G) Time spent in the EM versus S1 chamber during sociability. (H) Social preference index during sociability. Positive values indicate preference for S1. *n* = 11–12 mice per group. (I) Strategy for localized BCAA depletion in mPFC by accelerating catabolism. Schematic of BCAA catabolism and regulation of the BCKDH complex by BCKDK, with representative immunoblot for phosphorylated BCKDH E1α (p-E1α) and total E1α from control and mPFC *Bckdk* conditional knockout (*Bckdk*^ΔmPFC^) tissue. (J) Quantification of p-E1α/E1α ratio. *n* = 4 biologically independent samples per group. (K) Representative coronal section showing mPFC viral targeting and the track of the intracranial cannula used for local infusion (dashed outline indicates mPFC). Scale bar, 100 μm. (L) Targeted quantification of valine, leucine and isoleucine in mPFC tissue from control and *Bckdk*^ΔmPFC^ mice, normalized to control. *n* = 4 biologically independent samples per group. (M) Volcano plot of mPFC metabolomics comparing *Bckdk*^ΔmPFC^ versus control. Features meeting predefined thresholds (|log2 fold change| ≥ 0.6 and –log10 *P* > 1.3; dashed lines) are highlighted; BCAAs are annotated. (N) KEGG pathway enrichment analysis of differential metabolites from mPFC metabolomics. (O) Timeline for behavioural testing of *Bckdk*^ΔmPFC^ mice after viral injection. (P) Time spent in EM versus S1 during sociability in control and *Bckdk*^ΔmPFC^ mice. (Q) Social preference index during sociability in control and *Bckdk*^ΔmPFC^ mice. *n* = 17 and 19 mice per group, as shown. (R) Acute rescue paradigm. Vehicle or individual BCAAs (valine, leucine or isoleucine) were delivered into mPFC of *Bckdk*^ΔmPFC^ mice 30 min before sociability testing. (S) Time spent in EM versus S1 during sociability after acute intra-mPFC infusion. (T) Social preference index after acute intra-mPFC infusion, showing selective rescue by valine. *n* = 10–12 mice per group. Data are mean ± s.e.m. Dots represent individual mice unless otherwise indicated. In (G), (P), and (S), EM versus S1 was compared within each group by two-sided paired Student’s *t* tests. Group comparisons and multiple-testing procedures are described in the Methods. \**P* < 0.05, \*\**P* < 0.01, \*\*\**P* < 0.001; N.S., not significant.

Systemic restriction of essential amino acids, however, introduces physiological changes that complicate circuit-level interpretation. Across BCAA-restricted diets, mice exhibited reduced body weight (Figure S1B) and broad remodeling of CSF amino acid profiles beyond the targeted BCAA (Figure S1C). Open-field locomotor and center-exploration measures were largely preserved (Figures S1D–S1G), but gait analysis revealed abnormalities under a subset of diet conditions (Figures S1H–S1J). These systemic effects motivated a localized approach to reduce BCAA availability specifically within mPFC, a core hub for social behavior,^20–25^ while minimizing developmental and peripheral confounds.

To accelerate BCAA catabolism locally, we generated *Bckdk^ΔmPFC^* mice by bilaterally injecting AAV-Syn-Cre-EGFP into mPFC of *Bckdk^flox/flox^* mice;^28,29^ control littermates received AAV-Syn-EGFP (Figures 1I and S2A). Viral targeting was restricted to mPFC across animals (Figure S2A). Immunoblotting revealed reduced phosphorylation of the BCKDH E1α subunit (p-E1α), consistent with de-repression of BCKDH activity and enhanced local BCAA oxidation (Figures 1I and 1J). Targeted metabolomics confirmed reduced abundance of all three BCAAs in mPFC tissue, with valine exhibiting the largest depletion (Figure 1L). Untargeted metabolomics further revealed broad prefrontal remodeling, with BCAA-associated pathways among the most enriched signatures (Figures 1M and 1N). Importantly, *Bckdk^ΔmPFC^*mice displayed normal baseline locomotion and anxiety-like behavior in open field and O-maze assays (Figures S2B–S2F), preserved novelty-driven behavior in object recognition (Figure S2G), and largely intact levels of several non-BCAA essential amino acids in mPFC (Figure S2H), supporting a circuit-selective phenotype rather than generalized sickness, arousal, or global cognitive impairment.

Under these conditions of localized metabolic perturbation, *Bckdk^ΔmPFC^*mice displayed robust social impairment. Three weeks after viral injection, control mice showed strong sociability (S1 > EM), whereas *Bckdk^ΔmPFC^* mice failed to exhibit a significant preference for the social chamber (Figures 1O–1Q). A direct social interaction paradigm yielded a similar result: control mice habituated upon re-exposure and increased interaction with a novel conspecific, whereas *Bckdk^ΔmPFC^* mice lacked clear trial-dependent modulation (Figures S2I–S2K). Thus, local disruption of BCAA catabolic control in mPFC is sufficient to impair sociability, even in the absence of the systemic growth and motor confounds introduced by dietary restriction.

To determine which BCAA is behaviorally instructive within mPFC, we acutely supplemented individual BCAAs locally. Vehicle or valine, leucine, or isoleucine was infused into mPFC of *Bckdk^ΔmPFC^* mice 30 min before sociability testing (Figures 1R–1T), with cannula placement verified histologically (Figure 1K). Vehicle-treated *Bckdk^ΔmPFC^* mice remained asocial. In contrast, valine infusion robustly restored sociability, reinstating preference for S1 and rescuing the preference index (Figures 1S and 1T). Leucine or isoleucine produced at most modest and variable improvements that did not reliably restore the preference index to the same extent (Figures 1S and 1T). Together, the systemic discovery and mPFC-local rescue experiments identify valine as the uniquely instructive BCAA for mPFC-dependent sociability and indicate that its relevant action can occur on rapid, circuit-relevant timescales.

### LiP-SMap nominates mGluR5 as a valine-responsive synaptic effector

The rapid behavioral rescue produced by acute intra-mPFC valine suggested that valine is decoded by a signaling mechanism that can act acutely, rather than only by long-term nutritional or developmental effects. To identify candidate effectors without pre-specifying targets, we applied LiP-small molecule mapping (LiP-SMap)^30^ to native mPFC proteome extracts. In this approach, ligand engagement is inferred from ligand-dependent shifts in protease susceptibility under native conditions, detected as changes in the abundance of structure-informative conformotypic peptides by quantitative mass spectrometry (Figure 2A). LiP-SMap identified a set of valine-responsive features that passed stringent abundance and significance thresholds (Figure 2B). Among these, mGluR5 (Grm5) was supported by a valine-responsive peptide whose abundance increased upon valine exposure (Figure 2C), consistent with a ligand-dependent structural perturbation.

**Figure 2.**
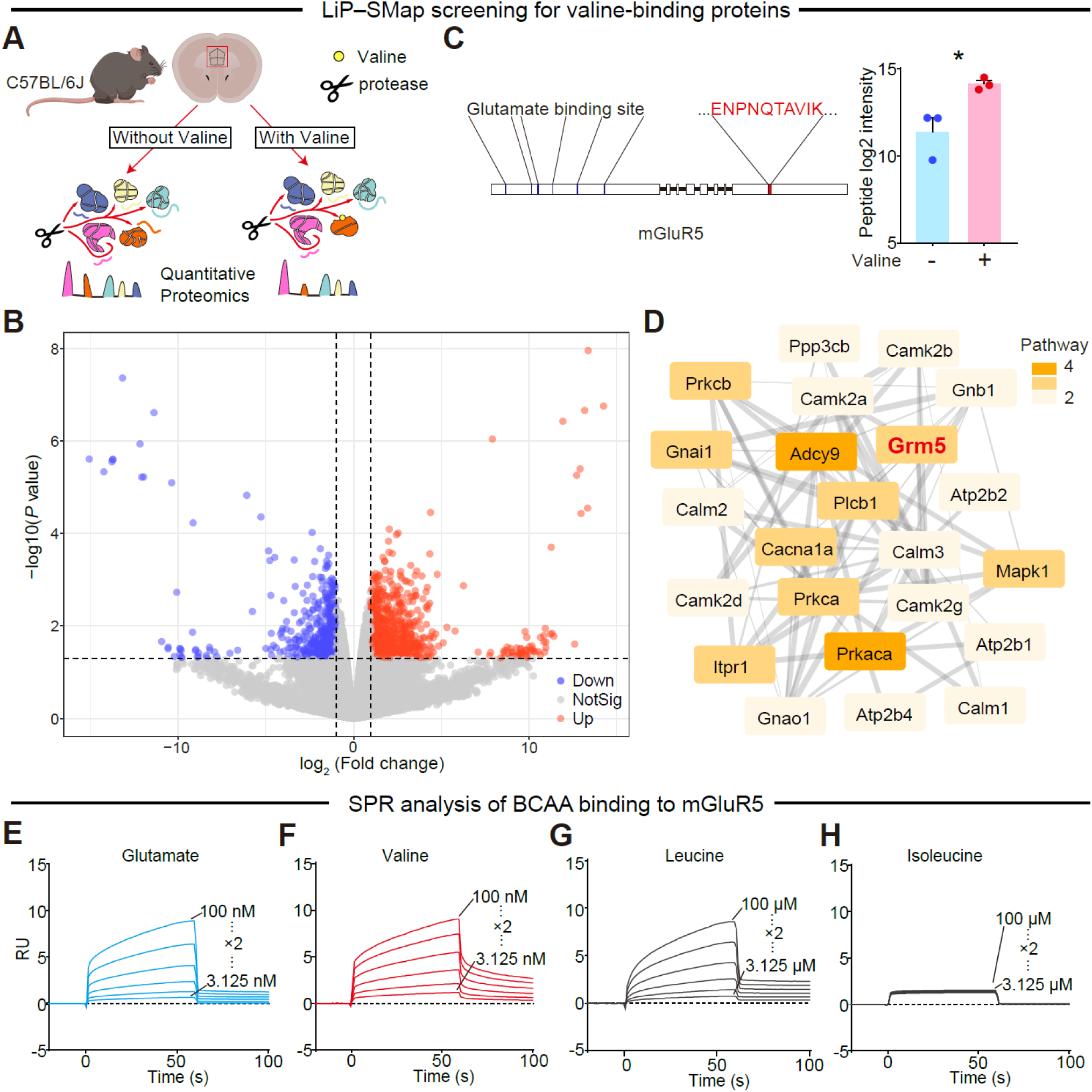
LiP-SMap and SPR identify mGluR5 as a valine-binding receptor. (A) Schematic of LiP-small molecule mapping (LiP-SMap). Native proteome extracts prepared from mouse mPFC were incubated with or without valine, subjected to limited proteolysis under native conditions, and processed for bottom-up quantitative mass spectrometry to identify ligand-dependent shifts in protease susceptibility (conformotypic peptides). (B) Volcano plot of LiP-SMap features comparing valine-treated versus untreated mPFC extracts. Dashed lines indicate predefined thresholds (|log_2_ fold change| ≥ 1 and –log_10_ *P* > 1.3). (C) A representative conformotypic peptide mapped to mGluR5 (left) and its abundance change upon valine treatment (right). The valine-responsive peptide (ENPNQTAVIK) lies outside the canonical glutamate-binding pocket region, consistent with a distinct structural perturbation. *n* = 3 biologically independent proteome preparations per condition. (D) Protein–protein interaction network of valine-responsive candidates integrated across four neurofunction-relevant pathways. Node shading indicates pathway breadth and edge thickness reflects interaction confidence. *Grm5* emerges as a multi-pathway hub. (E–h) Surface plasmon resonance (SPR) sensorgrams showing binding of glutamate (E), valine (F), leucine (G) and isoleucine (H) to mGluR5. Concentration series are indicated on the plots. In **c**, data are mean ± s.e.m. and dots represent independent proteome preparations. Significance was assessed by two-sided Welch’s *t* test. \**P* < 0.05.

To prioritize candidates capable of linking amino acid availability to synaptic computation and behavior, we integrated valine-responsive candidates into pathway-resolved interaction maps. This analysis highlighted four neurofunction-relevant programs—glutamatergic synapse, cAMP signaling, retrograde endocannabinoid signaling, and calcium signaling—that shared multiple nodes, suggesting convergence on a limited set of hubs (Figures 2D and S3A–S3D). Literature-based curation further emphasized Grm5 among these multi-pathway nodes, given its established roles in synaptic signaling and social-behavior-relevant phenotypes (Figure S3E). We therefore prioritized mGluR5 as a candidate receptor-level effector through which valine might tune prefrontal circuit function.

We next tested whether valine directly engages mGluR5 using surface plasmon resonance (SPR).^31^ SPR sensorgrams revealed robust binding of valine to mGluR5 in the nanomolar range, comparable in magnitude to glutamate, whereas leucine bound only weakly and isoleucine showed no detectable interaction under the same conditions (Figures 2E–2H). Receptor subtype profiling demonstrated selectivity: valine engagement of the closely related group I receptor mGluR1 was markedly weaker (Figure S4). These measurements validate mGluR5 as a selective BCAA-binding receptor and identify valine as the dominant mGluR5-engaging BCAA among those tested.

### Valine selectively binds mGluR5 and potentiates glutamate-evoked signaling

To define the functional consequence of valine binding, we first used PRESTO-Tango,^32^ a β-arrestin recruitment assay that couples arrestin engagement to transcriptional reporter output (Figure 3A). In this assay, both glutamate and valine produced dose-dependent increases in reporter signal, whereas leucine and isoleucine elicited comparatively weak responses (Figure 3B). Pharmacological dissection supported a non-orthosteric mechanism: the competitive orthosteric antagonist MCPG strongly reduced glutamate-driven reporter activation but failed to suppress valine-driven reporter output (Figures 3C and 3D), consistent with valine engaging mGluR5 through a mode distinct from canonical glutamate-site competition.

**Figure 3.**
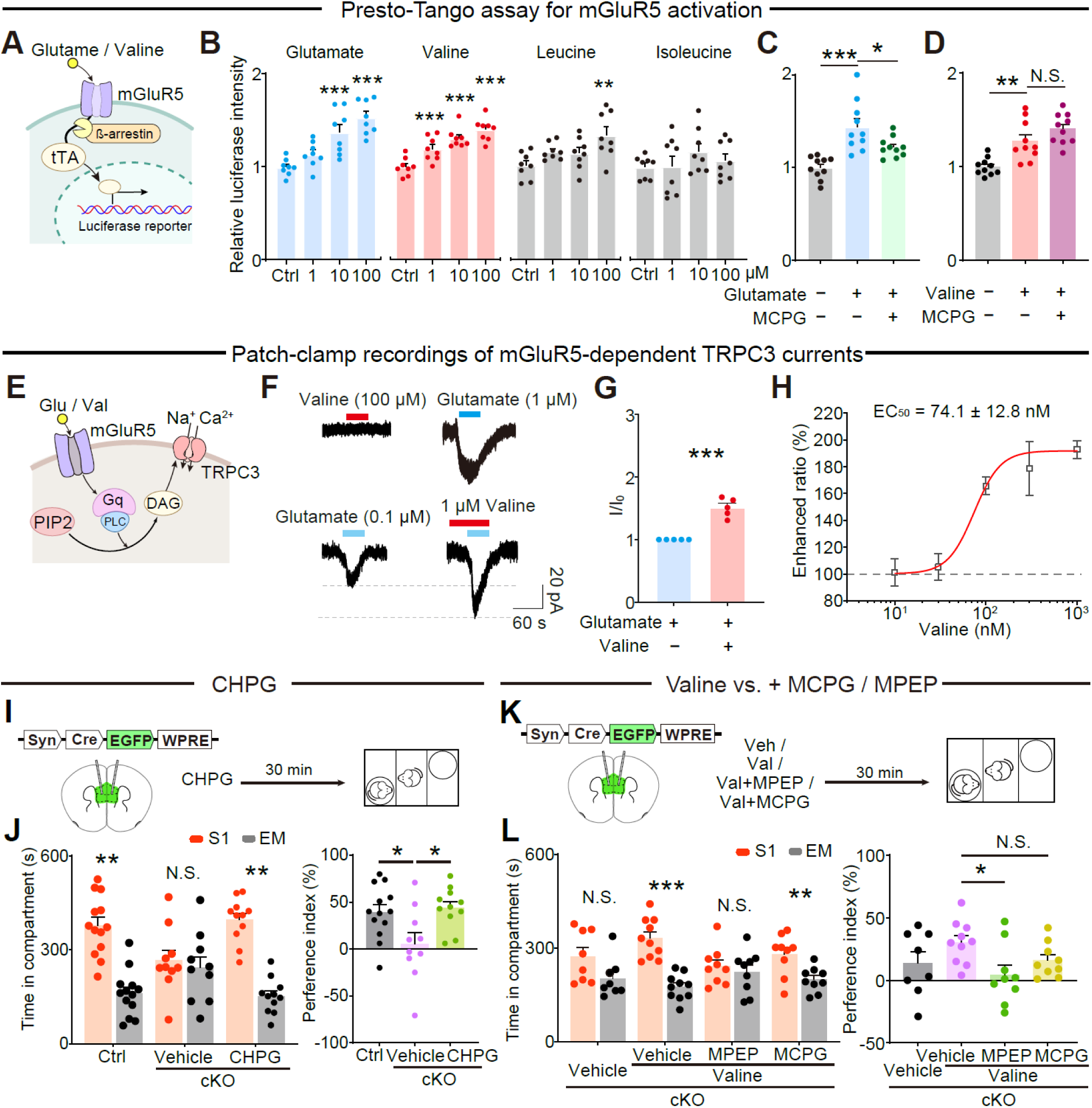
Valine potentiates mGluR5 through a non-orthosteric mechanism and rescues social deficits. (A) Schematic of the PRESTO-Tango β-arrestin recruitment assay used to quantify mGluR5 activation through transcriptional reporter output. Ligand-induced β-arrestin recruitment drives TEV-dependent release of a tethered transcription factor, leading to luciferase reporter expression. (B) PRESTO-Tango responses evoked by glutamate, valine, leucine and isoleucine at the indicated concentrations (0, 1, 10 and 100 μM). Dots indicate independent wells or measurements; *n* = 8 per condition. (C and D) Effects of the competitive orthosteric mGluR antagonist MCPG on PRESTO-Tango responses driven by glutamate (C) or valine (D). *n* = 9–10 per condition. (E) Schematic of the electrophysiological assay used to report canonical mGluR5–Gq–PLC signaling through TRPC3 currents in a heterologous system. (F) Representative whole-cell recordings. Top, 100 μM valine does not evoke detectable TRPC3 current, whereas 1 μM glutamate evokes a robust inward current. Bottom, pre-application of 1 μM valine enhances the response to subthreshold glutamate (0.1 μM). (G) Quantification of valine-dependent potentiation of subthreshold glutamate-evoked TRPC3 currents, expressed as a normalized current ratio (*I*/*I*_0_; see Methods). *n* = 5 cells per group. (H) Concentration–response relationship for valine-dependent potentiation, with EC_50_ indicated. (I) Experimental design for behavioural rescue with intra-mPFC infusion of the mGluR5 agonist CHPG before three-chamber testing in control and *Bckdk*^ΔmPFC^ mice. (J) Three-chamber performance shown as time spent in the social chamber (S1) versus the empty chamber (EM) and as social preference index. *n* = 13, 10, and 11 mice per group, as shown. (K) Experimental design for testing pharmacological constraints on valine-mediated rescue. *Bckdk*^ΔmPFC^ mice received vehicle or valine, alone or together with the mGluR5 negative allosteric modulator MPEP or the competitive mGluR antagonist MCPG, before the sociability assay. (L) Three-chamber performance for the conditions in (K). *n* = 8–10 mice per group, as shown. Data are mean ± s.e.m. In (B–D), statistics were analysed by one-way ANOVA with multiple comparisons as indicated. In **g**, a two-sided unpaired Student’s *t* test was used. In (J) and (L), EM versus S1 was compared within each group by two-sided paired tests and preference indices were compared between groups by two-sided unpaired tests, as indicated. \**P* < 0.05, \*\**P* < 0.01, \*\*\**P* < 0.001; N.S., not significant.

We next asked whether valine engages the canonical Gq arm of mGluR5 signaling. To obtain a temporally precise electrophysiological readout of receptor-driven Gq signaling, we co-expressed mGluR5 with the receptor-operated cation channel TRPC3 in a heterologous system and recorded TRPC3 inward currents, leveraging the well-established coupling of TRPC3 activation to GPCR–Gq pathways (Figure 3E).^33–36^ Whereas glutamate elicited robust inward currents, valine alone failed to evoke detectable TRPC3 currents under the same recording conditions (Figure 3F), indicating that valine is not an efficacious standalone agonist for this Gq-coupled output. By contrast, valine potentiated glutamate-evoked TRPC3 currents: valine pre-application amplified subsequent responses to subthreshold glutamate (Figures 3F and 3G) in a concentration-dependent manner with nanomolar potency (EC_50_ = 74.1 ± 12.8 nM; Figure 3H). To test whether this effect arises from modulation of TRPC3 itself rather than upstream receptor signaling, we activated endogenous Gq signaling independently of mGluR5 using the muscarinic agonist carbachol (CCH), which drives the mAChR–Gq–PLC pathway and in turn TRPC3 currents. Under these conditions, valine did not enhance CCH-evoked TRPC3 currents, indicating that valine acts upstream of the DAG–TRPC3 signaling module rather than on the module itself (Figure S5A). This potentiation was selective within the BCAA class: leucine did not enhance glutamate-evoked currents (Figures S5B and S5C). Isoleucine produced a potentiation-like effect in this assay (Figures S5D and S5E) despite lacking detectable mGluR5 binding by SPR (Figure 2H and Figure S4E), warranting cautious interpretation and leaving open mGluR5-independent contributions or context-dependent effects on the signaling module measured here.

If valine acts through mGluR5 *in vivo*, direct receptor activation should bypass valine deficiency. Consistent with this, intra-mPFC infusion of the mGluR5 agonist CHPG restored sociability in *Bckdk*^ΔmPFC^ mice, reinstating preference for the social chamber and rescuing the preference index (Figures 3I and 3J). Moreover, valine-mediated rescue was abolished by the mGluR5 negative allosteric modulator MPEP but persisted in the presence of MCPG (Figures 3K and 3L), matching the non-orthosteric pharmacology observed *in vitro*.

Together, these chemoproteomic, biophysical, and functional data support a model in which valine directly engages mGluR5 through a non-orthosteric mechanism and biases receptor output— supporting β-arrestin–linked signaling while acting as a potent positive allosteric modulator of glutamate-evoked canonical Gq signaling.

### A valine-selective VFT allosteric pocket underlies mGluR5 potentiation

We next asked where valine binds on mGluR5 and how this interaction potentiates receptor activation. To define the structural basis of valine engagement, we determined cryo-EM structures of full-length human mGluR5 under two ligand conditions: valine alone and valine plus glutamate (Figures 4, S6, and S7). From independent datasets processed with global refinement and focused local refinement of the extracellular Venus flytrap (VFT) domain, we obtained reconstructions at 3.86 Å (valine condition) and 3.19 Å (glutamate+valine condition), with VFT-local refinements reaching 3.33 Å and 2.89 Å, respectively, enabling residue-level analysis of ligand pockets (Figure S6).

**Figure 4.**
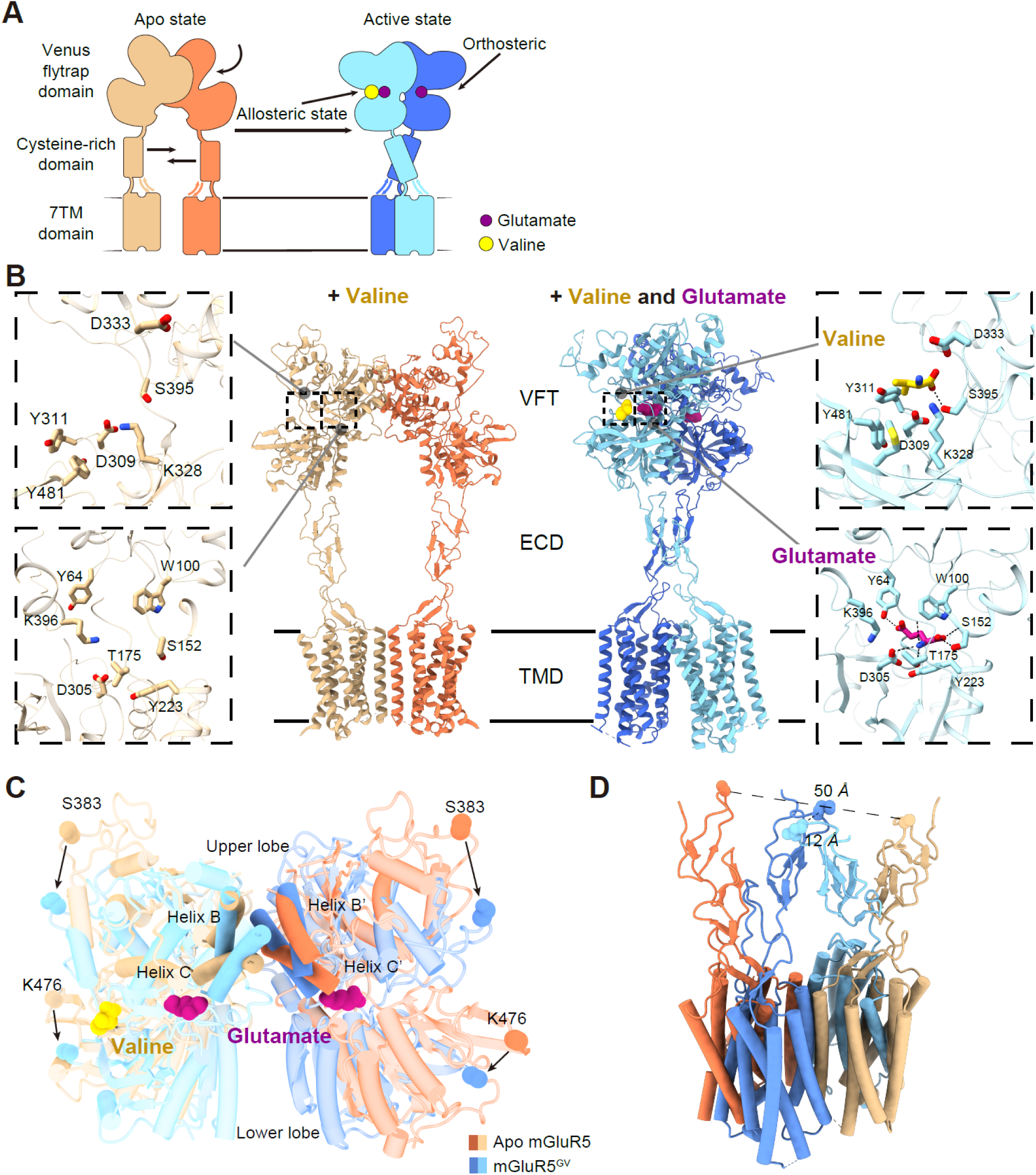
Cryo-EM reveals a valine-selective allosteric pocket in the mGluR5 Venus flytrap domain. (A) Domain organization and activation model of dimeric mGluR5, showing the Venus flytrap (VFT) domain, cysteine-rich domain (CRD) and seven-transmembrane domain (7TM/TMD). Glutamate binds the canonical orthosteric pocket in the VFT, whereas valine occupies a distinct allosteric pocket in the VFT. (B) Cryo-electron microscopy models of full-length human mGluR5 obtained in the presence of valine alone (left; apo-like conformation) or valine plus glutamate (right; active-like conformation). Insets show close-up views of the valine pocket (upper) and the orthosteric glutamate pocket (lower). Key side chains are shown as sticks; dashed lines indicate putative hydrogen bonds. (C) Superposition of VFT dimers from apo mGluR5 and glutamate-plus-valine-bound mGluR5 (mGluR5^GV^) highlighting ligand-coupled lobe closure and inter-protomer rearrangement. Upper-lobe movement is illustrated by displacement of S383, whereas K476 marks minimal movement in the lower lobe. Glutamate (magenta) and valine (yellow) are shown in their respective pockets. (D) CRD separation measured between the two protomers in apo mGluR5 (∼50 Å) and mGluR5^GV^ (∼12 Å), illustrating compaction of the extracellular assembly during activation.

The glutamate+valine structure (mGluR5^GV^) adopts an active-like architecture. Relative to the valine condition, the extracellular assembly is compacted, with pronounced approximation of the two cysteine-rich domains (CRDs) (Figures 4C and 4D), together with a corresponding reorganization of the transmembrane dimer (Figure S7E) consistent with class C GPCR activation.^37,38^ In the VFT, the orthosteric pocket contained clear density for glutamate, coordinated by canonical residues (Figures 4B and S7C). Strikingly, the VFT-local map also revealed a second, spatially distinct density consistent with valine at a previously unrecognized site adjacent to—but separable from—the orthosteric glutamate pocket (Figures 4B and S7C). This valine site resides at the upper/lower lobe interface of the VFT and presents a mixed polar–hydrophobic environment (Figure S7D). The pocket is lined by residues including K328, D333, S395, M469, Y480, and Y481, which together form a cavity suited to accommodate the valine side chain while positioning polar groups to engage the amino-acid backbone (Figures 4B and S7C). By contrast, under the valine-only condition we captured an apo-like conformation and did not observe comparably well-resolved density attributable to valine within the VFT, consistent with the functional observation that valine alone shows limited efficacy for canonical Gq/TRPC3 signaling while exerting its strongest effects in a glutamate-permissive context (Figures 3F–3H).

To test whether this structurally defined pocket is required for valine potentiation, we performed structure-guided mutagenesis of residues lining the pocket and quantified valine-dependent potentiation using the TRPC3 current assay as a direct readout of canonical mGluR5 signaling. In wild-type mGluR5, valine robustly enhanced glutamate-evoked inward currents (Figure 5A). Mutation of K328 (K328A), positioned to contribute to valine coordination, selectively abolished valine potentiation while preserving glutamate-evoked activation (Figure 5B), indicating that K328 is a critical determinant of valine allostery rather than general receptor competence. Extending this analysis, alanine substitutions at D333, S395, M469, Y480, and Y481 markedly reduced the potentiation index, whereas substitutions at nearby residues outside the pocket (for example, D309A and Y311A) did not significantly impair potentiation (Figure 5C). Thus, the same residues that define the valine pocket structurally are required for valine-dependent enhancement of glutamate-evoked signaling.

**Figure 5.**
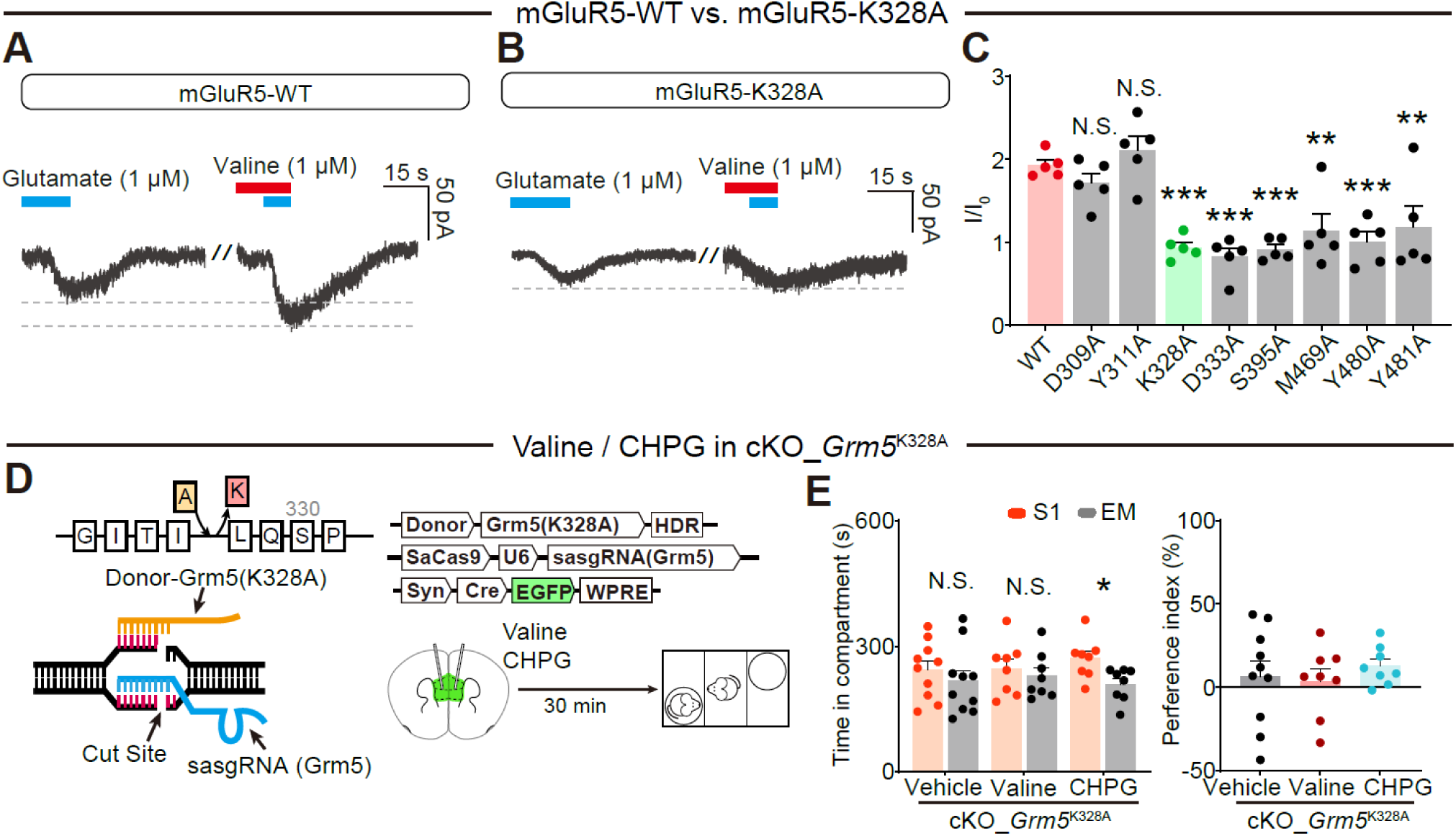
A single residue in the VFT pocket controls valine sensitivity of mGluR5. (A) Representative whole-cell recordings of mGluR5-dependent TRPC3 currents showing glutamate-evoked inward current and potentiation by valine in mGluR5 wild type (WT). (B) Representative recordings from the valine-pocket mutant mGluR5-K328A, showing preserved glutamate-evoked current but loss of valine-dependent potentiation. (C) Summary of valine-dependent potentiation across mGluR5 mutants surrounding the valine pocket, expressed as potentiation index (*I*/*I*_0_). Dots represent individual cells; *n* = 5 cells per group. (D) Schematic of CRISPR–Cas9-mediated introduction of the homologous K3278 substitution into mouse *Grm5* in mPFC, together with the behavioural testing paradigm. (E) Three-chamber performance in *Bckdk–Grm5(K328A)*^ΔmPFC^ mice after vehicle, valine or CHPG. Left, time spent in S1 versus EM; right, social preference index. *n* = 10, 8, and 8 mice per group, as shown. Data are mean ± s.e.m. In (C), multiple-group comparisons were analysed by one-way ANOVA with multiple comparisons as indicated. In (E), EM versus S1 was compared within each group by two-sided paired tests and preference index was compared between groups by two-sided unpaired tests, as indicated. \**P* < 0.05, \*\**P* < 0.01, \*\*\**P* < 0.001; N.S., not significant.

We next asked whether this pocket is required for valine-dependent behavioral rescue *in vivo*. To do this, we combined two manipulations in the same animals: Cre-mediated deletion of *Bckdk* in the mPFC of *Bckdk*^flox/flox^ mice to reduce local BCAA availability, and CRISPR–Cas9 editing of endogenous *Grm5* at the mouse residue homologous to human K328 (K327 in mouse; hereafter, the K328-equivalent site) (Figures 5D, S8A, and S8B). This design allowed us to test whether valine rescue depends specifically on the allosteric pocket, rather than on general receptor function, in the same local metabolic context used throughout the study. Sanger sequencing confirmed editing at the targeted *Grm5* locus. As shown before, intra-mPFC valine restored sociability in *Bckdk*^ΔmPFC^ mice (Figures 3K and 3L). By contrast, in *Bckdk*–*Grm5*(K328A)^ΔmPFC^ mice, valine failed to rescue the abnormal social phenotype (Figures 5E). Importantly, intra-mPFC administration of CHPG still restored social behavior in these mice (Figure 5E), indicating that the K328-equivalent site is specifically required for valine-dependent modulation *in vivo* but is dispensable for agonist-driven receptor activation *per se*.

Together, the structural, electrophysiological, and *in vivo* genetic data define a valine-selective allosteric pocket in the mGluR5 VFT domain that is distinct from the orthosteric glutamate site yet mechanistically coupled to glutamate-driven activation. Disrupting this pocket abolishes valine-dependent potentiation *in vitro* and prevents valine-mediated behavioral rescue *in vivo*.

### Valine-linked mGluR5–Gq signaling normalizes synaptic drive and inhibitory-network recruitment to preserve sociability

Having established a receptor-level mechanism, we next asked how impaired valine–mGluR5 coupling is expressed in mPFC physiology and whether restoring mGluR5/Gq output is sufficient to normalize circuit signatures and behavior in *Bckdk^ΔmPFC^* mice. We first examined baseline synaptic drive in acute mPFC slices. *Ex vivo* whole-cell recordings revealed that cKO neurons exhibit elevated spontaneous synaptic event rates. Relative to controls, cKO neurons showed increased sEPSC frequency with largely preserved sEPSC amplitude (Figures 6A–6C), as well as increased sIPSC frequency with minimal changes in sIPSC amplitude (Figures 6D–6F), indicating a frequency-dominant shift in spontaneous transmission. Acute engagement of mGluR5 with the selective agonist CHPG reduced the elevated sEPSC and sIPSC frequencies in cKO slices, shifting distributions toward lower event rates and decreasing per-cell mean frequencies with comparatively modest effects on amplitudes (Figures 6A–6F). These synaptic measurements indicate that local *Bckdk* loss is accompanied by heightened spontaneous synaptic event rates and that boosting mGluR5 activity is sufficient to counteract this signature.

**Figure 6.**
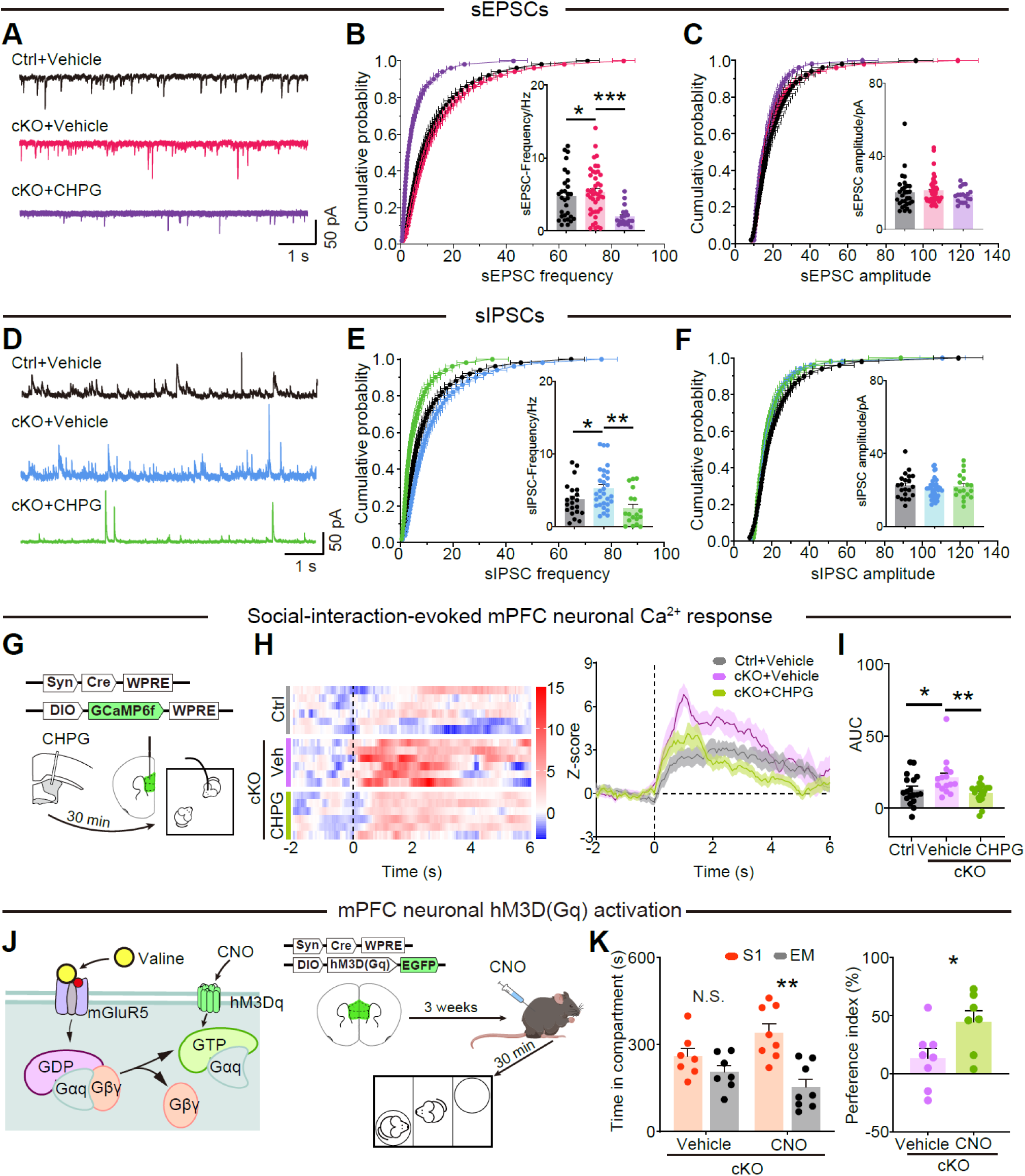
mGluR5–Gq signaling normalizes prefrontal circuit activity in *Bckdk*^ΔmPFC^ mice. (A) Representative whole-cell voltage-clamp traces of spontaneous excitatory postsynaptic currents (sEPSCs) recorded ex vivo from acute mPFC slices under the indicated conditions. (B) Cumulative probability distributions of sEPSC frequency. Inset, per-cell mean sEPSC frequency. (C) Cumulative probability distributions of sEPSC amplitude. Inset, per-cell mean sEPSC amplitude. *n* = 18-23 cells from 6–7 mice per group. (D) Representative whole-cell voltage-clamp traces of spontaneous inhibitory postsynaptic currents (sIPSCs) recorded ex vivo from acute mPFC slices under the indicated conditions. (E) Cumulative probability distributions of sIPSC frequency. Inset, per-cell mean sIPSC frequency. (F) Cumulative probability distributions of sIPSC amplitude. Inset, per-cell mean sIPSC amplitude. *N* = 18-32 cells from 6–7 mice per group. (G) Experimental design for mPFC fiber photometry during social interaction in control and *Bckdk*^ΔmPFC^ mice, with local CHPG or vehicle infusion before testing. (H) Social interaction-aligned photometry responses shown as trial heat maps (left) and group-averaged traces (right). Time 0 denotes onset of social investigation. (I) Quantification of photometry response magnitude as area under the curve (AUC) over the analysis window shown in (H). Each dot represents one social investigation bout; 15–18 bouts pooled from 5–6 mice per group. (J) Experimental design for chemogenetic bypass of receptor activation using mPFC expression of hM3Dq(Gq) in *Bckdk*^ΔmPFC^ mice and acute CNO administration before behavioural testing. (K) Three-chamber performance after vehicle or CNO in *Bckdk*^ΔmPFC^ mice. Left, time spent in S1 versus EM; right, social preference index. *n* = 7–8 mice per group. Data are mean ± s.e.m. In (B), (C), (E), (F), and (I), multiple-group comparisons were analysed by one-way ANOVA with multiple comparisons as indicated. In (K), EM versus S1 was compared within each group by two-sided paired tests and preference index was compared between groups by two-sided unpaired tests. \**P* < 0.05, \*\**P* < 0.01, \*\*\**P* < 0.001; N.S., not significant.

We next asked whether this synaptic phenotype is mirrored by altered circuit recruitment during social behavior. Fiber photometry recordings of mPFC population calcium signals during social investigation revealed an exaggerated social-evoked response in cKO mice relative to controls, quantified as increased response magnitude (AUC) over the interaction-aligned epoch (Figures 6G–6I). Local CHPG delivery suppressed this heightened response toward control levels (Figures 6H and 6I), linking mGluR5 engagement to normalization of a behavioral-state-dependent network signature. To test whether downstream signaling is sufficient to bypass impaired metabolic coupling, we expressed hM3Dq(Gq) in mPFC and acutely activated Gq signaling with CNO. Chemogenetic Gq activation improved sociability and increased the preference index in cKO mice (Figures 6J and 6K), supporting the mGluR5–Gq axis as a sufficient entry point to recover social approach under conditions of disrupted local BCAA/valine availability.

### Valine–mGluR5 engages endocannabinoid/CB1 feedback to gate inhibitory transmission during social interaction

The CHPG-sensitive suppression of social-evoked hyperactivity suggested that valine–mGluR5 coupling constrains recruitment of a specific circuit component during social engagement. We therefore performed cell-type-biased calcium recordings in mPFC during social interaction. Excitatory neuron-enriched photometry (CaMKIIα-GCaMP6f) did not reveal a significant cKO-associated increase in the social-evoked response and was not detectably altered by CHPG at the AUC level (Figure S9). By contrast, inhibitory neuron-enriched photometry (mDlx-GCaMP6f) revealed a pronounced cKO-associated increase in social-evoked activity that was significantly reduced by CHPG (Figure S9). These data identify excessive inhibitory-network recruitment as a prominent signature of the cKO social state and show that mGluR5 activation constrains this hyper-response during social engagement.

Because this inhibitory-network phenotype appeared disproportionate to the cellular distribution of mGluR5, we next assessed the expression context of *Grm5* in mPFC. RNAscope co-localization indicated that *Grm5* is detected in a larger fraction of *Vglut1*^+^ excitatory neurons than *Vgat*^+^ inhibitory neurons (36.60% versus 12.90% *Grm5*^⁺^; Figure S10), arguing against a simple explanation in which inhibitory selectivity arises from preferential receptor expression in inhibitory neurons. This mismatch suggested that valine–mGluR5 might act through a trans-synaptic mechanism that biases impact toward inhibitory circuitry.

One established route by which postsynaptic mGluR5 can impose presynaptic control is retrograde endocannabinoid signaling, in which receptor engagement promotes endocannabinoid release that acts on presynaptic CB1 receptors to suppress transmitter release.^39–43^ Guided by this framework, we mapped the anatomical substrate for CB1-mediated gating in mPFC. RNAscope for *Cnr1* revealed strong enrichment in inhibitory neurons: ∼60.5% of *Vgat*^+^ cells were *Cnr1*^+^, compared with ∼12.3% of *Vglut1*^+^ cells (Figures 7A and 7B). This distribution predicts that an mGluR5-triggered retrograde signal would act with disproportionate efficacy at inhibitory synapses.

**Figure 7.**
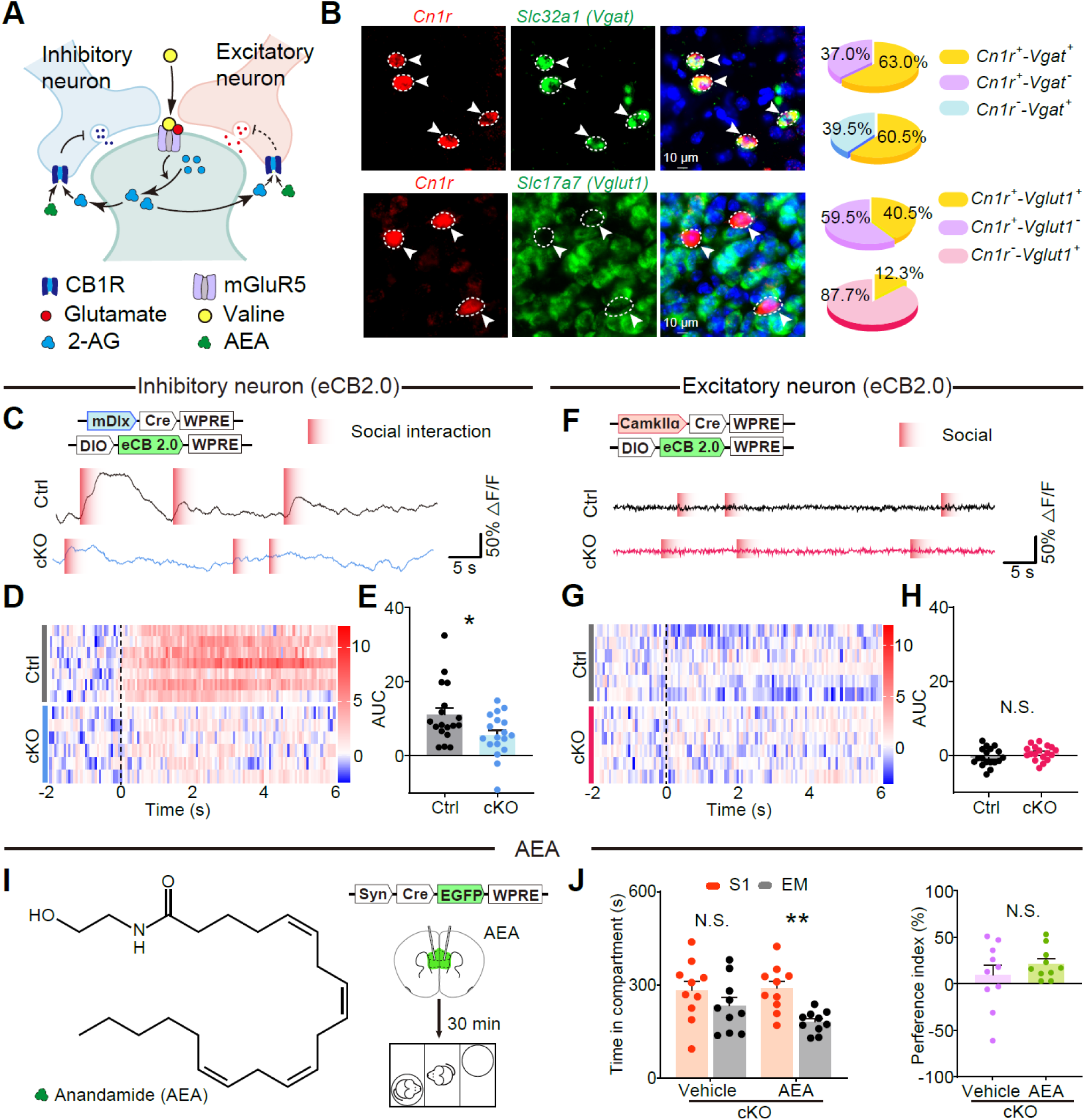
Endocannabinoid–CB1 signaling links valine–mGluR5 coupling to inhibitory gating. (A) Working model. Valine allosterically enhances glutamate-dependent mGluR5 signaling to promote endocannabinoid release, which acts retrogradely on presynaptic CB1 receptors to suppress transmitter release. Enrichment of CB1 on inhibitory terminals biases this feedback towards functional disinhibition. (B) RNAscope/ISH analysis of *Cnr1* (CB1 receptor; red) co-localized with inhibitory (*Slc32a1/Vgat*, green; top) or excitatory (*Slc17a7/Vglut1*, green; bottom) markers in mPFC. Pie charts report the fraction of *Cnr1*^+^ cells that are *Vgat*^+^ or *Vglut1*^+^, and conversely the fraction of *Vgat*^+^ or *Vglut1*^+^ cells that are *Cnr1*^+^. Scale bars, 10 μm. *n* = 3 mice per group, with 3 sections analysed per mouse. (C) Cell-type-restricted monitoring of endocannabinoid dynamics in inhibitory neurons using the genetically encoded sensor eCB2.0. Representative *ΔF/F* traces are shown for control and *Bckdk*^ΔmPFC^ mice; red-shaded periods indicate social investigation. (D) Heat maps of z-scored eCB2.0 signals aligned to the onset of social investigation (t = 0; dashed line) for inhibitory-neuron recordings. (E) Quantification of inhibitory-neuron eCB2.0 responses as AUC over the analysis window shown in (D). Each dot represents one social investigation bout; 18 bouts pooled from 6 mice per group. (F) Parallel strategy and representative traces for eCB2.0 monitoring in excitatory neurons. (G) Heat maps of z-scored eCB2.0 signals aligned to social onset for excitatory-neuron recordings. (H) Quantification of excitatory-neuron eCB2.0 responses as AUC. Each dot represents one social investigation bout; 18 bouts pooled from 6 mice per group. (I) Chemical structure of the CB1 agonist anandamide (AEA) (left) and experimental design for intra-mPFC AEA or vehicle administration before sociability testing in *Bckdk*^ΔmPFC^ mice (right). (J) Three-chamber performance after intra-mPFC vehicle or AEA in *Bckdk*^ΔmPFC^ mice. Left, time spent in S1 versus EM; right, social preference index. *n* = 10 mice per group. Data are mean ± s.e.m. In (E) and (H), groups were compared using two-sided unpaired Student’s *t* test. In (J), EM versus S1 was compared within each group by two-sided paired tests and preference index was compared between conditions by two-sided unpaired tests, as indicated. \**P* < 0.05, \*\**P* < 0.01; N.S., not significant.

To test whether endocannabinoid signaling is altered during social engagement, we monitored endocannabinoid dynamics *in vivo* using the genetically encoded sensor eCB2.0 targeted to defined neuronal populations.^44,45^ When eCB2.0 was expressed in mPFC inhibitory neurons (mDlx-Cre), social investigation in control mice was accompanied by a robust, time-locked increase in sensor signal, whereas cKO mice showed a significantly blunted response, quantified as reduced AUC (Figures 7C–7E). In contrast, when eCB2.0 was targeted to excitatory neurons (CaMKIIα-Cre), socially aligned sensor responses were weaker and more variable and did not differ significantly between control and cKO mice (Figures 7F–7H). Thus, loss of local BCAA/valine homeostasis is associated with a selective reduction in social-state endocannabinoid signaling at inhibitory circuit elements, consistent with diminished CB1 engagement at inhibitory synapses.

Finally, we asked whether restoring CB1 signaling is sufficient to improve sociability. Intra-mPFC infusion of the CB1 agonist anandamide (AEA) reinstated a significant within-group preference for the social chamber (S1 > EM) in cKO mice, although the preference index did not differ significantly between vehicle and AEA groups (Figures 7I and 7J), indicating a partial rescue under these conditions. Taken together with the CHPG-sensitive normalization of inhibitory-network recruitment and social behavior (Figures 6 and S9), these data support a circuit mechanism in which valine amplifies glutamate-dependent mGluR5 signaling to promote endocannabinoid/CB1 feedback that preferentially gates inhibitory transmission during social interaction, thereby constraining inhibitory-network over-recruitment and preserving sociability.

## Discussion

This study identifies a direct route by which amino acid availability can shape cortical circuit function. The central finding is that valine is not simply one component of a broader BCAA state, nor only a substrate for protein synthesis and energy metabolism. In mPFC, valine is decoded by a valine-selective allosteric site on mGluR5, where it biases receptor output and increases the gain of glutamate-evoked activation. *In vivo*, this coupling engages endocannabinoid/CB1 signaling to gate inhibitory transmission, constrain inhibitory-network recruitment during social interaction, and preserve mPFC-dependent sociability. Two points are important at the outset. *First*, the relevant variable is valine specificity, not generic BCAA tone. *Second*, the bridge from metabolic state to social behavior is implemented not only through slower nutrient-sensing or proteostatic programs, but through direct engagement of a neurotransmitter receptor, a design that is well suited to circuit timescales.

### Valine functions as a synaptic signal beyond nutrient supply

Valine is usually discussed as a substrate. It supports protein synthesis, contributes to energy metabolism through BCAA oxidation, and participates in broader nutrient-sensing networks.^11,12,19,46^ Yet work in multiple systems suggests that valine abundance can also function as a signal—a variable that is detected and translated into specific cellular outcomes. In hematopoiesis, changing valine availability reshapes stem and progenitor compartments.^47^ In cancer and metabolic settings, valine handling can influence mitochondrial and stress programs through translation-linked mechanisms.^48^ More recently, direct valine-binding proteins have been identified that use valine abundance to regulate defined outputs.^49^ In parallel, BCAA catabolism generates acyl-CoA intermediates that feed into protein acylation,^50^ providing a slower route by which valine flux may affect cellular function. Together, these examples suggest that valine can be more than permissive—it can be interpreted.

Our findings extend this idea to the nervous system in a mechanistically distinct way. Canonical intracellular amino acid sensors, such as mTORC1 or integrated stress-response pathways, primarily read out cytosolic pools and typically act over minutes to hours, with broad effects on translation and cellular adaptation.^19^ Here, valine influences behavior through a receptor at the cell surface, providing a route by which valine can tune synaptic signaling with behavioral timing. This distinction suggests at least two coupled layers of valine biology in brain: intracellular valine, which supports biosynthesis and nutrient sensing, and extracellular valine, which can tune receptor function. Both are likely to matter in neurodevelopmental conditions,^15–18^ but they differ in compartment, kinetics, and potentially in therapeutic accessibility.

### Valine–mGluR5 allostery converts tonic availability into glutamate-contingent gain

A defining feature of the mechanism is glutamate contingency. In our assays, valine supported β-arrestin-linked reporter output yet showed little standalone efficacy for canonical Gq/TRPC3 coupling, while it strongly potentiated glutamate-evoked mGluR5 signaling. This profile makes sense in the context of synaptic physiology. Glutamate release is phasic and tightly coupled to activity,^26^ whereas extracellular valine is likely to vary more slowly because it is shaped by diet, BBB, uptake, and local metabolism. An allosteric mechanism that scales glutamate-driven activation therefore provides a straightforward way to translate a comparatively tonic metabolic variable into phasic modulation that remains aligned with synaptic timing. In other words, valine does not impose tonic receptor activation; it changes how strongly mGluR5 responds when glutamate is present.

The structural data sharpen this logic. Cryo-EM and pocket mutagenesis define a valine-selective VFT allosteric pocket that is distinct from the orthosteric glutamate site and required for potentiation. The glutamate+valine condition captures an active-like architecture, whereas valine alone does not stabilize the same activation signature, consistent with the functional finding that valine is most effective in a glutamate-permissive context. A simple interpretation is that valine shifts the ensemble of glutamate-bound receptor states toward conformations that couple more efficiently to downstream signaling. This is compatible with the view that class C GPCR activation proceeds through multiple steps and that different ligands can bias state occupancy along that path.

The structural data are consistent with this view. Valine binds a VFT allosteric pocket that is distinct from the orthosteric glutamate site and is required for potentiation. The glutamate-plus-valine condition captures an active-like receptor architecture, whereas valine alone does not stabilize the same activation signature. One interpretation is that valine shifts the glutamate-bound ensemble toward states that couple more efficiently to downstream signaling. This fits well with the idea that class C GPCR activation proceeds through multiple steps and that orthosteric ligands and allosteric modulators stabilize different parts of that pathway.^37,38,51–53^ In this framework, valine is best understood as a physiological allosteric ligand with biased output. This also places limits on what can be concluded. Our data establish receptor selectivity within group I mGluRs, because valine binds and functionally modulates mGluR5 much more effectively than mGluR1. They do not establish whether valine influences other glutamatergic receptors, including AMPA and NMDA receptors, either directly or indirectly.^27,54,55^ We did not find evidence pointing to ionotropic receptors as primary valine effectors in the present study, but whether valine can affect them remains an important open question.

Because the valine pocket is extracellular, the relevant ligand for mGluR5 is extracellular valine in the interstitial or synaptic space, yet the perturbation in *Bckdk^ΔmPFC^*mice is intracellular and metabolic: accelerated BCAA catabolism lowers tissue valine, with valine being depleted most strongly. This means that any receptor-level effect must be coupled, directly or indirectly, to changes in extracellular valine near synapses. One plausible route is transport-driven pool coupling. If intracellular valine becomes limiting, neurons may increase uptake from extracellular space, lowering local extracellular valine and reducing occupancy of the mGluR5 pocket. Alternatively, altered exchange through amino acid transporters could remodel extracellular composition without requiring large changes in total tissue abundance.^56^ These local dynamics must also integrate BBB transport of large neutral amino acids and neuron–glia exchange.^17^ The acute rescue produced by local valine delivery is consistent with extracellular sufficiency, but the quantitative relation between intracellular consumption, extracellular microdomains, and receptor occupancy remains unknown. Direct measurements of interstitial valine during behavior, coupled to manipulations of transport pathways, will be needed to resolve this point.

### Valine–mGluR5 recruits endocannabinoid/CB1 feedback to gate inhibition in mPFC

At the circuit level, the clearest signature of impaired valine coupling is state-dependent over-recruitment of inhibitory networks during social interaction, while excitatory-neuron population signals are comparatively preserved. This is not explained simply by receptor distribution, because *Grm5* is detected in a larger fraction of excitatory than inhibitory neurons in mPFC. The data instead point to a trans-synaptic implementation, in which postsynaptic mGluR5 signaling is converted into a presynaptic control signal that lands preferentially on inhibitory circuitry.

Retrograde endocannabinoid signaling provides such a mechanism.^40–42,57^ In many circuits, group I mGluRs engage Gq/PLCβ-linked pathways that mobilize endocannabinoids, which then act through presynaptic CB1 receptors to suppress transmitter release.^39^ A recurring principle, especially clear in hippocampus, is that CB1 receptors are enriched on selected inhibitory terminals, allowing endocannabinoids to regulate inhibition with high specificity.^57^ Our data fit this logic closely. In mPFC, *Cnr1* expression is biased toward inhibitory neurons, and endocannabinoid sensor signals during social engagement are selectively blunted in inhibitory circuitry when valine availability is reduced. This provides a direct explanation for why mGluR5 activation, or downstream Gq engagement, normalizes inhibitory-network recruitment and restores sociability. It also helps explain why elevated spontaneous inhibitory event rates can coexist with exaggerated population activity during social interaction: if mGluR5–endocannabinoid feedback is weakened, inhibitory networks may be recruited excessively or inappropriately rather than being properly gated by behavioral state.

The downstream arm of the pathway is likely broader than the endocannabinoid branch alone. In this study, TRPC3 served as a readout of canonical mGluR5–Gq–PLC signaling in a heterologous system. But TRPC3 is not merely an artificial reporter; it is expressed in central neurons and has been implicated in metabotropic synaptic responses.^58^ This raises the possibility that valine-dependent tuning of mGluR5 could affect not only endocannabinoid release but also other native outputs downstream of Gq/PLC, including Ca^2+^ dynamics, slow depolarizing conductances, and forms of synaptic plasticity. The divergence between β-arrestin-linked reporter output and canonical Gq/TRPC3 signaling also suggests that valine may not amplify all mGluR5 pathways equally. Determining which downstream branches are most sensitive to valine occupancy will be important both for physiology and for any future attempt to manipulate this site therapeutically.

### Implications for neurodevelopmental dysfunction

BCAA metabolism and transport have been linked to neurodevelopmental phenotypes through metabolomics, human genetics (including *BCKDK*), and BBB transport mechanisms. mGluR5 has also been implicated in synaptic and behavioral abnormalities relevant to autism and related conditions,^59–63^ although the direction of dysfunction varies by context. One implication of our data is that mGluR5 function can be weakened without mutation in the receptor itself, simply through loss of an endogenous allosteric input that normally amplifies glutamate-driven signaling. This may help explain why mGluR5-directed interventions have not had uniform effects across disease models: the critical question may not be whether mGluR5 activity is globally high or low, but whether circuits are operating with the correct activity-contingent gain.

From a translational perspective, the valine pocket offers a structural template for glutamate-contingent modulation, which in principle could preserve physiological timing and avoid some of the liabilities of tonic receptor activation. At the same time, any approach based on valine or valine-pocket agonism will need to account for valine’s essential intracellular roles and for the likelihood that chronic manipulation of valine handling will engage additional pathways beyond mGluR5.

### Limitations of the study

Several limitations remain clear. We have not measured extracellular valine dynamics in mPFC during behavior, which is essential for linking metabolism to receptor occupancy *in vivo*. The identity and source of the endocannabinoid signal remain unresolved, as does the necessity of CB1 signaling in specific inhibitory microcircuits for the behavioral phenotype. We also do not yet know whether sustained manipulation of valine–mGluR5 coupling reshapes synaptic plasticity and network stability over longer timescales, or how general the mechanism is across cortical regions and developmental stages. Finally, although our data establish receptor selectivity within group I mGluRs, the broader question of whether valine can modulate ionotropic glutamate receptors remains open.

## Conclusion

In summary, valine acts as a diet-derived synaptic signal in mPFC by engaging a valine-selective allosteric site on mGluR5. This converts amino acid availability into glutamate-contingent receptor gain and recruits endocannabinoid/CB1 feedback to gate inhibition during social behavior. This pathway provides a direct molecular route from amino acid availability to cortical circuit control of sociability and offers a framework for understanding metabolic–synaptic coupling in neurodevelopmental dysfunction.

## STAR★METHODS

Detailed methods are provided in the online version of this paper and include the following:

**KEY RESOURCES TABLE**

**RESOURCE AVAILABILITY**

Lead contact

Materials availability

Data availability

Code availability

## EXPERIMENTAL MODEL AND SUBJECT DETAILS

Mice

## METHOD DETAILS

Stereotaxic surgery and implantation

Viral vectors and local pharmacology

Behavioral assays

CSF and mPFC tissue collection

Targeted and untargeted metabolomics

Immunoblotting

Recombinant mGluR5 expression and purification

Cryo-EM sample preparation and data collection

Cryo-EM image processing

Model building and refinement

LiP-small molecule mapping (LiP-SMap)

Pathway and protein-interaction analysis

Surface plasmon resonance

PRESTO-Tango assay

Cell culture and electrophysiology

CRISPR–Cas9 editing of the *Grm5* valine-pocket residue *in vivo*

Acute slice electrophysiology

Fiber photometry and signal processing eCB2.0 sensor recordings

Immunofluorescence

RNAscope *in situ* hybridization

Image analysis

**QUANTIFICATION AND STATISTICAL ANALYSIS**

**SUPPLEMENTAL INFORMATION**

Supplemental Information includes 10 Figures and 3 Tables, and their legends are available for this paper online.

## ACKNOWLEDGMENTS

This study was supported by grants from the National Natural Science Foundation of China (82430104, 82125032, 82301729, 82404755, 22206128, and 82204048), the China Postdoctoral Science Foundation (2022M712125), the Science and Technology Commission of Shanghai Municipality (YDZX20253100003001, 23Y21900500, 23DZ2291100, and 2018SHZDZX01), the Shanghai Municipal Health Commission (GWV-11.1-34, GWVI-11.2-YQ30, 2020CXJQ01, and 2018YJRC03), the Innovative Research Team of High-level Local Universities in Shanghai (SHSMU-ZDCX20211100), the Shanghai Oriental Talents Program, the Chenguang Program of Shanghai Education Development Foundation and Shanghai Municipal Education Commission, and “Discipline Peak-Climbing Plan” of Xinhua Hospital Affiliated to Shanghai Jiao Tong University School of Medicine (XKPF2024A50011).

## AUTHOR CONTRIBUTIONS

X.S., T.Z., J.-Y.Z., W.-G.L., and F.L. conceived the project, designed the experiments, and interpreted the results. X.S. and T.Z. performed most of the behavioral experiments, animal surgery, Ca^2+^ recording, eCB sensor recording, immunohistochemistry, RNA scope staining, and data analysis. H.L., Yuhua Yan, Y.G., and L.S. assisted with some of the behavioral experiments and conducted viral injections. S.W., K.H., and D.W. performed PRESTO-Tango assay. X.S., T.Z., Z.-Y.Z., H.-R.G., and J.Y. performed LiP-SMap and SPR. X.Z. and Ye Yu performed cellular electrophysiology. T.-T.S. performed slice electrophysiology. Y.W., X.Y., and Q.Q. performed structural determination. X.S., Y.G., and Q.Z. performed metabolomics analysis. W.Z., L.Z., and B.M. commented on data interpretation. H.S. generated *Bckdk*^flox/flox^ mice. X.S., T.Z., X.W., W.-G.L., and F.L. wrote the manuscript with contributions from all authors. All authors read and approved the final manuscript.

## DECLARATION OF INTERESTS

The authors declare no competing interests.

## INCLUSION AND DIVERSITY

We support inclusive, diverse, and equitable conduct of research.

## KEY RESOURCES TABLE

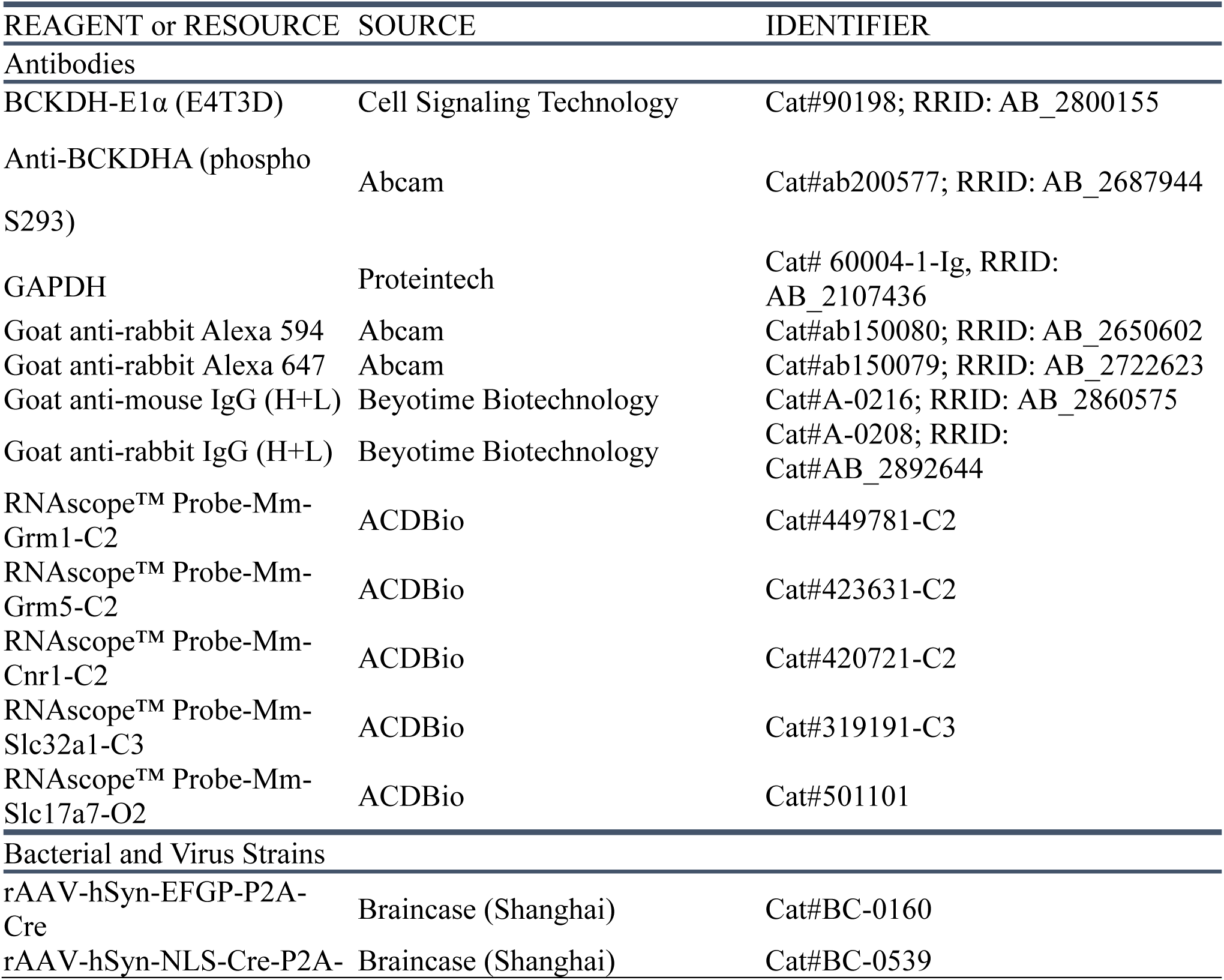
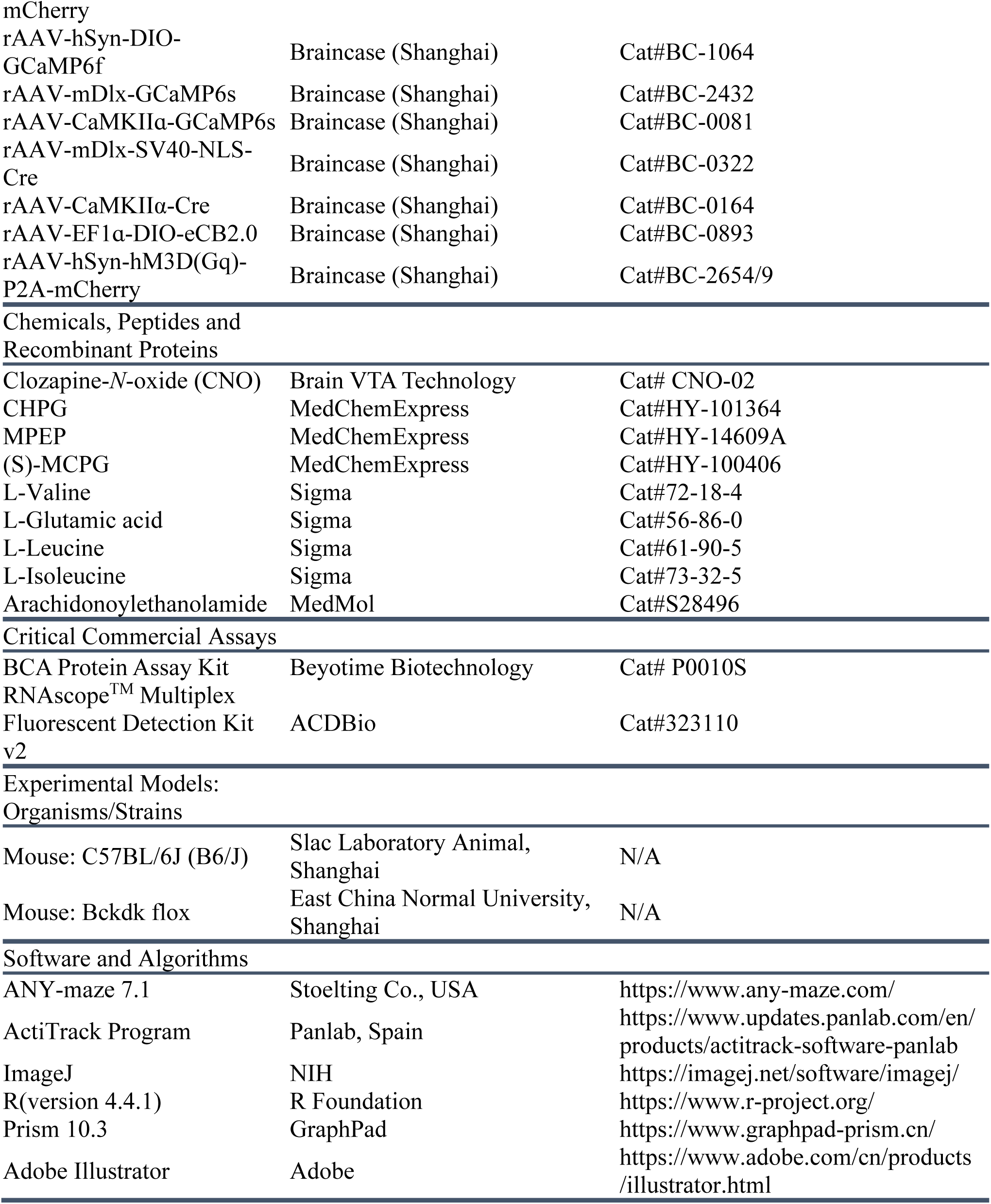

## RESOURCE AVAILABILITY

### Lead contact

Further information and any related requests should be directed to and will be fulfilled by the lead contact, Prof. Fei Li. All published reagents can be shared on an unrestricted basis.

## Materials availability

This study did not generate new unique reagents.

## Data availability

All data supporting the findings of this study are available within the paper, the Supplementary Information and the associated Source Data files. Cryo-EM data collection and refinement statistics are provided in Extended Data Table 1. The coordinates for Apo mGluR5 and mGluR5GV have been deposited in the Protein Data Bank (PDB) under accession codes 20YN and 9XFO, respectively. The cryo-EM density maps for Apo mGluR5 and mGluR5GV have been deposited in the Electron Microscopy Data Bank (EMDB) with accession codes EMD-67397 and EMD-66836, respectively. Additional data are available from the corresponding author upon reasonable request.

## Code availability

No custom code central to the main conclusions of this study was generated. Data acquisition and analysis used commercial or publicly available software, including ANY-maze, ActiTrack, cryoSPARC, RELION, GraphPad Prism, MATLAB, Clampex and Clampfit. Any custom scripts used for data organization, plotting or secondary analysis are available from the corresponding author upon reasonable request.

## EXPERIMENTAL MODEL AND SUBJECT DETAILS

### Mice

Wild-type C57BL/6J mice were purchased from Shanghai SLAC Laboratory Animal Co., Ltd., Shanghai, China. *Bckdk*^flox/flox^ mice on a C57BL/6J background were generated as previously reported.^28,29^ Mice were maintained in a specific pathogen-free animal facility at the Institute of Brain Functional Genes, East China Normal University, under a 12-h light/12-h dark cycle at 22–25 °C and 55–60% humidity, with *ad libitum* access to standard rodent chow and water. Unless otherwise indicated, male mice were used for behavioral experiments. Mice were group-housed (up to five animals per cage) before experiments. All procedures were approved by the Animal Ethics Committee of East China Normal University (Ethics approval number: m20220104), and every effort was made to minimize animal numbers and suffering.

## METHOD DETAILS

### Stereotaxic surgery and implantation

For viral delivery, 4-week-old mice were anaesthetized with 1.5–2.0% isoflurane (R510-22, RWD Life Science) in 100% oxygen and positioned in a stereotaxic frame (RWD Life Science, China). The skull was levelled by aligning bregma and lambda in the same horizontal plane. Viral vectors were delivered bilaterally into mPFC at the following coordinates relative to bregma: anterior–posterior (AP), +1.75 mm; medial–lateral (ML), ±0.35 mm; dorsal–ventral (DV), −2.60 mm. Virus was infused at 0.1 μl min^−1^ using a Hamilton microsyringe, with a total volume of 0.35 μl per hemisphere. The needle was left in place for 10 min after infusion and then withdrawn slowly.

Mice recovered in their home cages for at least 3 weeks to allow robust viral expression. For fiber-photometry experiments, an optic fiber assembly (200-μm core, 1.25-mm ferrule, 3-mm fiber length) was implanted 0.02 mm above the viral injection site (AP, +1.75 mm; ML, ±0.35 mm; DV, −2.58 mm). For local pharmacology, guide cannulae were implanted at a 30° angle to avoid the sagittal sinus and target mPFC at AP, +1.70 mm; ML, ±1.65 mm; DV, −1.05 mm. Implants were secured to the skull using stainless-steel screws and dental acrylic. Mice recovered for at least 1 week before behavioral testing. Viral expression and placement of optic fibers or cannulae were verified post hoc by fluorescence imaging.

### Viral vectors and local pharmacology

AAV vectors used in this study included AAV-Syn-Cre-EGFP (Braincase, cat. no. BC-0160) and AAV-Syn-EGFP (Braincase, cat. no. BC-0020) for local manipulation in *Bckdk*^flox/flox^ mice, CaMKIIα-GCaMP6f (BrainVTA, cat. no. PT-0119) and mDlx-GCaMP6f (BrainVTA, cat. no. PT-3025) for cell-type-biased fiber photometry, Cre-dependent hM3Dq (Gq) (Braincase, cat. no. BC-1246) for chemogenetic activation of Gq signaling, and Cre-dependent eCB2.0 (Braincase, cat. no. BC-0893) for monitoring endocannabinoid dynamics in defined neuronal populations. Viral delivery was performed as described above.

For local pharmacological manipulations, compounds were delivered bilaterally into mPFC through guide cannulae. Valine (Sigma-Aldrich, cat. no. V0513), leucine (Sigma-Aldrich, cat. no. L8912), isoleucine (Sigma-Aldrich, cat. no. I2752), CHPG (MedChemExpress, cat. no. HY-101364), MPEP (MedChemExpress, cat. no. HY-14609A), MCPG (MedChemExpress, cat. no. HY-100371), AEA (MedChemExpress, cat. no. HY-10863) and the corresponding vehicle solutions were infused 30 min before behavioral testing unless otherwise indicated in the relevant figure legends. CNO (Braincase, cat. no. CNO-25mg) was administered before testing in hM3Dq experiments as indicated in the main text and figure legends.

### Behavioral assays

Behavioral testing was performed during the light phase. Before testing, mice were handled for 3 consecutive days (3–5 min per day) and acclimated to the testing room for 1 h. Apparatuses were cleaned with 75% ethanol between animals. Different behavioral assays were separated by at least 1 week.

#### Three-chamber sociability assay

Three-chamber sociability testing was performed with minor modifications from previous work.^64^ The apparatus (60 cm × 40 cm × 25 cm) consisted of three communicating chambers separated by Perspex walls with central openings (8 cm × 12 cm). Two identical wire cups were placed in the side chambers. Mice were first acclimated to the apparatus for 5 min and then allowed to habituate freely to all three chambers for 10 min. Time spent in each side chamber during habituation was recorded to exclude innate side preference.

For the sociability phase, an unfamiliar age- and sex-matched conspecific (stranger 1, S1), previously habituated to the wire cup, was placed in one side chamber, whereas the other side chamber contained an empty cup (EM). The location of S1 was counterbalanced across animals. The test mouse was then allowed to explore the apparatus freely for 10 min. Chamber occupancy and exploration were recorded using ANY-maze (Stoelting Co., USA). The social preference index was calculated as:

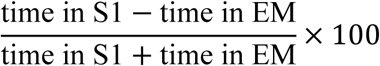

so that positive values indicate preference for S1, consistent with the figures.

#### Open field test

Locomotor activity and anxiety-like behavior were assessed in a square Plexiglas arena (25 cm × 25 cm × 30 cm). Mice were placed in the center of the arena and allowed to explore freely for 30 min. Activity was analysed using ActiTrack (Panlab, Spain), with distance travelled sampled in 5-min bins. Total distance travelled over 30 min was used as a measure of locomotion. Center distance, center time and center entries during the first 5 min were used as measures of anxiety-like behavior.

#### Elevated O-maze

Anxiety-like behavior was also assessed in an elevated O-maze consisting of two open and two enclosed quadrants. Each mouse was tested once for 5 min, starting from a closed quadrant, and behavior was recorded with ANY-maze (Stoelting Co., USA). Time spent in open arms, number of open-arm entries and distance travelled in open arms were quantified.

#### Novel object recognition

Novel object recognition was performed in the same open-field arena under dim illumination. During the familiarization phase, mice were allowed to explore two identical objects for 6 min. After a 2-min delay, one object was replaced by a novel object, and mice were returned to the arena for a 3-min test phase. Exploration time for the familiar and novel objects was quantified using ANY-maze. The discrimination index was calculated as:

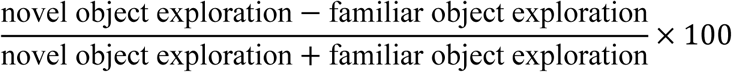

#### Direct social interaction assay

The direct social interaction assay was used to assess social interaction, habituation and social novelty in an unconstrained setting. Partner mice were partially shaved before testing to facilitate identification during scoring. Test mice were first acclimated to the recording environment for 15 min. A partner mouse was then introduced for a 2-min interaction period (T1), and interaction time was recorded. After a 30-min interval with the test mouse alone, the same partner mouse was reintroduced for a second 2-min interaction period (T2) to assess habituation. After another 30-min interval, a novel partner mouse was introduced for a third 2-min interaction period (T3) to assess social novelty.

#### Gait analysis

Gait was evaluated using a footprint assay. Mouse paws were coated with non-toxic ink and animals walked along a paper-lined corridor toward a dark goal box. Footprint patterns were collected from at least three successful runs per mouse, and gait length and gait width were manually measured from the footprints using ImageJ by an experimenter blinded to experimental conditions.

### CSF and mPFC tissue collection

#### CSF collection

Cerebrospinal fluid (CSF) was collected after the dietary manipulation period for metabolomic analyses. Mice were anaesthetized and positioned with the head flexed downward. CSF was withdrawn from the cisterna magna using a glass capillary under visual guidance. Samples visibly contaminated with blood were excluded. CSF was transferred immediately to low-bind tubes, snap-frozen on dry ice and stored at −80 °C until metabolomic analysis.

#### mPFC tissue collection

For metabolomics and immunoblotting, mice were deeply anaesthetized and rapidly decapitated.

Brains were removed into ice-cold buffer, and bilateral mPFC tissue was dissected on an ice-cold platform using the same anatomical boundaries as the viral targeting region. Samples were snap-frozen in liquid nitrogen and stored at −80 °C until use.

### Targeted and untargeted metabolomics

#### CSF metabolomics

CSF samples were thawed on ice and extracted with chilled organic solvent to precipitate proteins and recover metabolites. After vortexing and centrifugation, clarified supernatants were transferred to autosampler vials and analyzed by liquid chromatography–mass spectrometry (LC–MS)-based metabolomics. For targeted analysis, valine, leucine, and isoleucine were quantified against internal standards and expressed relative to the control group. For untargeted profiling, metabolite features were subjected to peak detection, alignment, normalization, and quality-control filtering before downstream analysis. Principal component analysis and row-scaled heat maps were generated from the processed feature matrix.

#### mPFC metabolomics

Frozen mPFC tissue was weighed and homogenized in ice-cold methanol: acetonitrile: water (2:2:1) containing internal standards. Homogenates were vortexed, incubated on ice, and centrifuged to remove debris, and the resulting supernatants were subjected to LC–MS analysis. Targeted quantification of valine, leucine, and isoleucine was performed using external calibration curves and normalized to tissue input. For untargeted metabolomics, LC–MS features were extracted and aligned across samples using Progenesis QI, normalized to internal standards, and corrected for signal drift using pooled quality-control samples prior to statistical analysis. Differential metabolites were visualized by volcano plot (|log_2_ fold change| ≥ 0.6 and –log_10_ *P* > 1.3), followed by KEGG pathway enrichment analysis.

### Immunoblotting

mPFC tissue was homogenized in ice-cold lysis buffer containing protease and phosphatase inhibitors. Lysates were clarified by centrifugation, and protein concentration was determined by BCA assay or equivalent. Equal amounts of protein were separated by SDS–PAGE and transferred to PVDF membranes. Membranes were blocked in 5% non-fat milk or bovine serum albumin in Tris-buffered saline with Tween-20 and incubated overnight at 4 °C with primary antibodies against phosphorylated BCKDH E1α (p-E1α) (Cell Signaling Technology, cat. no. 90198, 1:1000) and total BCKDH E1α (Abcam, cat. no. ab200577, 1:1000). After washing, membranes were incubated with HRP-conjugated secondary antibodies and developed using chemiluminescent substrate. Band intensities were quantified in ImageJ or equivalent software, and the p-E1α/E1α ratio was calculated for each sample. Antibody details should be listed in the Key Resources Table.

### Recombinant mGluR5 expression and purification

Human mGluR5 (residues 21–872) was cloned into a modified pEG BacMam expression vector.^65^ For cryo-electron microscopy, the N terminus was fused to a haemagglutinin signal peptide, a FLAG tag and a twin-Strep tag. Bacmids were generated in DH10Bac *Escherichia coli* and used to produce recombinant baculovirus in *Spodoptera frugiperda* Sf9 cells.

For expression, 1 L of Expi293 cells were cultured in Gibco^®^ FreeStyle^TM^ 293 Expression Medium (ThermoFisher Scientific) at 37 °C under a 5% CO_2_ atmosphere to reach a density of 3 × 10^6^ cells per ml. 50 ml of P3 viruses produced using above Bac-to-Bac system (Invitrogen) were added to the culture. After 12 h, the cell culture was supplemented with 10 mM sodium butyrate and the culture temperature was changed to 30 °C. After another 48–60 h, cells were harvested by centrifugation at 1,500 × g for 15 min, washed once with PBS, frozen in liquid nitrogen, and stored at −80 °C.

For purification, cell pellets collected from 1 L of culture were resuspended in buffer A (25 mM HEPES, 150 mM NaCl, 5% glycerol, pH 7.4) containing 1% *N*-Dodecyl-β-D-maltoside (DDM, Anatrace), 0.1% cholesteryl hemisuccinate (CHS, Anatrace), 1mM phenylmethylsulphonyl fluoride (PMSF), and 3 μg/mL protease inhibitor cocktail (Aprotinin: Pepstatin: Leupeptin (w/w) = 1:1:1). The resuspended cell pellets were dounced in a 50-mL glass homogenizer for 40 strokes on ice, followed by gentle agitation at 4 °C for 3 h. The cell debris was removed by centrifuging at 16,000 rpm at 4 °C for 30 min. The supernatant was incubated with 2 mL of pre-equilibrated Streptactin Beads 4FF (Smart-Lifesciences, cat no. SA053025) and stirred gently at 4 °C for 2 h. Beads were packed into a gravity column (Smart-Lifesciences, cat no. SLM001) and washed with 10 column volume (CV) of buffer A containing 0.1% DDM and 0.01% CHS before being incubated with 1% lauryl maltose neopentyl glycol (LMNG, Anatrace) and 0.1% CHS in buffer A at 4 °C for 1 h. The column was washed with 30 CV of buffer A containing LMNG and CHS at concentrations ranging from (0.1% LMNG + 0.01% CHS) to (0.001% LMNG + 0.0001% CHS). Recombinant mGluR5 was eluted with 5 mM D-Desthiobiotin (ChemCruz, cat. no. sc-294239A), 0.001% LMNG, and 0.0001% CHS in buffer A. The pooled fractions were concentrated with a 100 kDa cut-off concentrator (Merck Millipore, Cat no. UFC810096,) and then subjected to size-exclusion chromatography using a Superose 6 10/300 GL column (Cytiva, cat. no. 29-0915-96,) equilibrated with buffer B (25 mM HEPES, 150 mM NaCl, 0.001% LMNG, 0.0001% CHS, pH 7.4). The peak fractions corresponding to purified recombinant mGluR5 were concentrated to 7.25 mg/mL for cryo-EM grid preparation.

For samples supplemented solely with valine (1 mM valine added during protein purification), only the apo-state conformation of mGluR5 was successfully resolved. For mGluR5^GV^ (mGluR5 bound to glutamate and valine), 10 mM L-(+)-glutamic acid (Beyotime, cat no. 56-86-0,) and 10 mM L-(+)-valine (Sigma-Aldrich, cat. no. 72-18-4) were added throughout the protein purification process. The sample was aliquoted, flash-frozen in liquid nitrogen, and stored at −80 °C for further use.

### Cryo-EM sample preparation and data collection

Protein quality was first assessed by negative-stain electron microscopy. Purified mGluR5 was applied to glow-discharged copper grids coated with a thin carbon film, stained with 2% uranyl acetate and imaged on a Talos L120C transmission electron microscope operated at 120 kV.

Cryo-EM grids were prepared as described previously.^66^ Briefly, Quantifoil Au 1.2/1.3 (300 mesh) grids were glow-discharged (15 mA, 55 s), loaded with 2.3 μl purified protein, blotted for 3–5 s at 4 °C and 100% humidity in a Vitrobot Mark IV, and plunge-frozen into liquid ethane. Grids were stored in liquid nitrogen until data collection.

All datasets were collected on a Titan Krios G4 transmission electron microscope operated at 300 kV and equipped with a Falcon G4i direct electron detector and Selectris X energy filter (20-eV slit). Movies were acquired using EPU software in super-resolution mode at a calibrated pixel size of 0.932 Å and a defocus range of −1.2 to −2.0 μm. The total dose per electron event representation (EER) movie was approximately 50 e^−^/Å^2^.

### Cryo-EM image processing

All datasets were processed similarly using cryoSPARC (v.3.3.2)^67^ and RELION (v.3.1.4).^68^ The dataset of the mGluR5 sample in apo state included 5,298 EER movies. Each EER movie of 1080 frames was fractionated into 40 subgroups, and beam-induced motion was corrected with a MotionCor2-like algorithm implemented in RELION. Exposure-weighted micrographs were then imported into cryoSPARC for CTF (contrast transfer function) estimation by patch CTF. Particles were initially blob-picked and extracted with a box size of 360 pixels and subjected to multiple rounds of 2D classification. Several rounds of heterogeneous refinement (3D classification) were conducted using ab initio reference maps reconstructed with 2D averages of nice features. The good particles were then converted for Bayesian polishing in RELION, which was subsequently imported back to cryoSPARC for one more round of ab initio reconstruction and several rounds of heterogeneous refinement to remove residual contaminants or poor-quality particles. The final 3.86 Å Apo mGluR5 map was generated from 74,019 particles by non-uniform refinement. After local refinement with a VFT domain mask, the 3.33 Å Apo mGluR5_VFT_ map was obtained. Resolution of the map was estimated internally in cryoSPARC by gold-standard Fourier shell correlation with the 0.143 criterion. The glutamate–valine dataset (mGluR5^GV^) was processed in the same way. From 7,220 EER movies, a final set of 85,209 particles yielded a global mGluR5^GV^ map at 3.19 Å and a VFT-focused map at 2.89 Å.

### Model building and refinement

Initial models were based on inactive and active mGluR5 structures from Protein Data Bank entries 8T6J and 8TAO. Ligand models and geometry restraints were generated using eLBOW in Phenix.^69^ Models were manually adjusted in Coot^70^ and refined in real space with Phenix (v.1.19).^69^ Side chains lacking clear density were truncated before deposition. Model geometry was evaluated with MolProbity.^71^ Final refinement statistics are summarized in Supplemental Table 1.

### LiP-small molecule mapping (LiP-SMap)

LiP-small molecule mapping was performed essentially as described previously,^30^ with modifications for mouse mPFC extracts. Native protein extracts were prepared from mPFC tissue in a non-denaturing buffer compatible with limited proteolysis. Protein concentration was normalized across samples before incubation with valine or vehicle. After ligand incubation, samples were subjected to limited proteolysis under native conditions to generate conformation-sensitive fragments, and the reaction was then quenched and followed by denaturation and complete digestion for bottom-up proteomics. Peptides were analysed by quantitative LC–MS/MS. Ligand-responsive features were defined as peptides showing reproducible abundance changes between valine-treated and untreated conditions. Differential features were visualized by volcano plot, and peptides mapping to candidate proteins were integrated with pathway and protein-interaction analyses.

### Pathway and protein-interaction analysis

Proteins identified by LiP-SMap as valine-responsive candidates were mapped onto neurofunction-relevant pathways, including glutamatergic synapse, cAMP signaling, retrograde endocannabinoid signaling and calcium signaling, using curated pathway databases and interaction resources. Protein–protein interaction networks were visualized to identify nodes represented in multiple pathway modules. Literature-based annotation of selected candidate hubs included reported human genetic links, ASD-related phenotypes and SFARI evidence categories where applicable. Nodes represented across multiple pathway modules and supported by prior disease or circuit evidence were prioritized for further validation.

### Surface plasmon resonance

Direct binding of amino acids to recombinant human mGluR5 and mGluR1 was measured by surface plasmon resonance (SPR).^31^ Purified receptor proteins, in the assay format used experimentally, were immobilized on the sensor chip according to established procedures. Serial dilutions of glutamate, valine, leucine and isoleucine were injected over the chip surface, and association and dissociation were recorded in real time. Reference-subtracted sensorgrams were analysed using the instrument software or equivalent fitting procedures to obtain kinetic parameters and apparent equilibrium dissociation constants.

### PRESTO-Tango assay

PRESTO-Tango was performed with minor modifications from the published protocol.^32^ HTLA cells, a HEK293-derived cell line stably expressing a tTA-dependent luciferase reporter and a β-arrestin2–TEV fusion protein, were maintained in DMEM supplemented with 10% fetal bovine serum and penicillin–streptomycin (100 U/mL and 100 μg/mL, respectively) at 37 °C in a humidified atmosphere containing 5% CO₂. HTLA cells were transiently transfected with plasmids encoding the full-length human metabotropic glutamate receptor 5 (mGluR5, wild-type) constructed in the PRESTO-TANGO backbone using Lipo8000™ (Beyotime, cat. no. C0533-0.5ml), according to the manufacturer’s instructions. Twenty-four hours after transfection, cells were dissociated and replated into white, clear-bottom 96-well plates at a density of approximately 4 × 10⁴ cells per well in 100 μl of complete medium. For receptor activation, cells were incubated with compounds diluted in a custom branched-chain amino acid–free medium (BOSTER) for the indicated durations. Luciferase activity was measured using Bright-Glo™ Luciferase Assay System (Promega). Briefly, culture medium was removed, and 100 μl of 20-fold diluted Bright-Glo reagent prepared in HEPES-buffered HBSS (20 mM HEPES, pH 7.4) was added to each well. After incubation at room temperature for 15–20 min, luminescence was recorded using a microplate reader (Gen5). Luminescence signals were expressed as relative light units (RLU). Relative receptor activation was calculated by normalizing drug-treated signals to control wells after background subtraction. Data were analysed using GraphPad Prism. All experiments were performed in at least three independent biological replicates with multiple technical replicates per condition.

### Cell culture and electrophysiology

Human embryonic kidney (HEK-293) cells were cultured in DMEM medium adding 1% penicillin/streptomycin (Gibco), 1% glutamate (Gibco) and 10% FBS (PAM) at 37°C in a humidified atmosphere of 5% CO_2_ and 95% air. The plasmids of human mGluR5-EGFP and hTRPC3 was transiently co-transfected in HEK293 cells in a ratio 1:1 with the transfection agent containing two types of solutions (solution A contains 250 mM CaCl_2_ in pure water; solution B contains (in mM) 1.5 Na_2_HPO_4_, 140 NaCl, and 50 HEPES, and pH was adjusted to 6.96).

All electrophysiological recordings were performed at room temperature (23 ± 2℃) with Axon200B on HEK293 cells after at least 24 h transfection.^72^ The conventional whole-cell configuration was performed under the voltage clamp with a holding potential of –60mV. Patch pipettes had a resistance ranging from 3-5 MΩ when filled with intracellular solution containing (in mM) 30 NaCl, 0.5 CaCl_2_·2H_2_O, 10 HEPES, 120 KCl, 1 MgCl_2_·6H_2_O and 5 EGTA with pH adjusted to 7.4 by Tris-base.^73^ Throughout the whole-cell recordings, cells were bathed in the standard external solution containing (in mM): 150 NaCl, 5 KCl, 10 glucose, 2 CaCl_2_·2H_2_O, 10 HEPES and 1 MgCl_2_·6H_2_O with pH being adjusted to 7.4. All agonists were dissolved and diluted by the standard external solution. Membrane current signals were amplified by a patch clamp amplifier (Axon200B, Axon Instruments, Foster City, CA) with low-pass filtered at 1 kHz using low-pass Bessel filters and 0.5 of output gain (α) (low-pass filter at 2kHz and 50 of output gain (α) for single-channel recordings). A Digidata 1440B interface and a computer running the Clampex and Clampfit 10.6 software (Molecular Devices) were used to sample and analyze all currents. A “Y-tube” method was used to perfuse solutions and compounds throughout all electrophysiological recordings.

### CRISPR–Cas9 editing of the *Grm5* valine-pocket residue *in vivo*

To disrupt the valine-binding pocket of mGluR5 *in vivo*, the mouse residue homologous to human K328 was identified as K327 in *Grm5*. For clarity and to facilitate direct comparison with the human structure, this homologous mouse residue is referred to throughout the manuscript as the K328-equivalent site unless otherwise specified. A CRISPR–Cas9 strategy was designed to introduce the K327A substitution in mPFC. sgRNAs were selected to target the genomic region surrounding the K327 codon, and donor templates containing the desired substitution and flanking homology arms were prepared to support homology-directed repair. CRISPR–Cas9 reagents were delivered into mPFC using the expression system indicated in the main text and figure legends. After recovery, editing at the target locus was verified by PCR amplification of genomic DNA from the targeted region followed by Sanger sequencing.

### Acute slice electrophysiology

Whole-cell recordings were performed in acute brain slices from *Bckdk^flox/flox^*mice that had been stereotaxically injected with AAV-Syn-Cre-EGFP or AAV-Syn-EGFP in mPFC brain regions. Mice were deeply anesthetized with Zoletil 50 (50 mg/kg) and were subsequently decapitated. Brains were dissected quickly and were chilled in well-oxygenated (95% O_2_ / 5% CO_2_, v/v) ice-cold artificial cerebrospinal fluid (ACSF) containing the following (in mM): 125 NaCl, 2.5 KCl, 12.5 D-glucose, 1 MgCl_2_, 2 CaCl_2_, 1.25 NaH_2_PO_4_, and 25 NaHCO_3_ (pH 7.35–7.45), with a pH between 7.35 and 7.45. Coronal brain slices (300 μm thick) containing regions of mPFC were cut with a vibratome (Leica VT1200 S, Germany). After recovery for 40 min in oxygenated ACSF at 34 ± 1 °C, each slice was transferred to a recording chamber and was continuously superfused with oxygenated ACSF at a rate of 1–2 mL per minute. To assess the effects of Val and CHPG on fluorescently labelled neurons, Val (100 mM) and CHPG (200 nM) was added to the oxygenated ACSF respectively before each slice was superfused and incubate for another 30 min to ensure optimal cellular uptake of the drug and thereby facilitate its pharmacological activity. The neurons in the mPFC were patched under visual guidance using an infrared differential-interference contrast microscope (BX51WI, Olympus, Japan) and an optiMOS camera (QImaging, Teledyne Imaging Group, USA). The slices were continuously perfused with well-oxygenated ACSF at 34 ± 1°C during all electrophysiological studies. Whole-cell patch clamp recordings were performed using an Axon 200B amplifier (Molecular Devices, USA). Membranous currents were sampled and analysed using a Digidata 1550 interface and a personal computer running Clampex and Clampfit software (Version 10.5, Molecular Devices, USA). Access resistance was 15–30 MΩ and only cells with a change in access resistance < 20% were included in the analysis. The brain slice received illumination via a 40× water-immersion lens (LUMPLFLN 40XW, Olympus, Japan).

### Fiber photometry and signal processing

Fluorescence signals were acquired using a dual-wavelength fiber-photometry system (RWD Life Science, China) with 470-nm excitation for Ca^2+^-dependent or sensor-dependent fluorescence and 410-nm excitation as the isosbestic control. Excitation light was delivered through a patch cord connected to the implanted optic fiber via a rotary joint to permit unrestricted movement. Power at the fiber tip was adjusted to 20–30 μW before each session. Signals were sampled at 30–90 Hz and acquired using the RWD fiber-photometry software. Behavioral events were synchronized to fluorescence acquisition by video recording.

Before behavioral recording, mice were habituated to the patch cord and recording chamber to minimize stress and movement artefacts. Each mouse was placed in the recording chamber and acclimated for 5 min while tethered, followed by a 1-min fluorescence baseline. A novel conspecific was then introduced and fluorescence signals were recorded continuously for 5 min.

Raw photometry signals were processed using custom MATLAB workflows and/or equivalent analysis pipelines. The 410-nm control channel was used to correct for motion and bleaching, and corrected fluorescence was expressed as *ΔF/F*. For event-based analyses, traces were aligned to the onset of social investigation. Z-scoring was performed relative to a pre-event baseline window. Heat maps and group-averaged traces were generated from aligned trials, and event magnitude was quantified as the area under the curve over the analysis window defined in the corresponding figure.

### eCB2.0 sensor recordings

To monitor endocannabinoid signaling *in vivo*, the genetically encoded sensor eCB2.0 was expressed in mPFC neurons using cell-type-restricted strategies. For inhibitory-neuron-biased recordings, eCB2.0 was expressed using an mDlx-based strategy (Braincase, cat. no. BC-0322); excitatory-neuron-biased recordings used a CaMKIIα-based strategy (Braincase, cat. no. BC-0164). Fiber photometry was then performed as described above. Social investigation served as the event-aligned behavioral epoch. For each recording, fluorescence traces were synchronized to behavior and aligned to the onset of social investigation. Signal amplitude was quantified as area under the curve over the analysis window specified in the corresponding figure.

As eCB2.0 reports endocannabinoid-pathway engagement rather than the identity of a single endogenous ligand, interpretations in the manuscript are restricted to endocannabinoid signaling unless otherwise stated.

### Immunofluorescence

Adult C57BL/6J mice (8–12 weeks old) were deeply anaesthetized with pentobarbital and transcardially perfused with ice-cold PBS followed by 4% paraformaldehyde in PBS. Brains were removed, post-fixed overnight at 4 °C and sectioned coronally at 35 μm on a vibratome. Free-floating sections were washed in PBS, permeabilized and blocked in PBS containing 5% normal goat serum and 0.3% Triton X-100 for 1 h at room temperature.

Sections were incubated overnight at 4 °C with primary antibodies diluted in blocking solution, including anti-VGLUT1 (HUABIO, cat. no. HA601449, 1:500) and anti-VGAT (HUABIO, cat. no. HA601384, 1:500), as appropriate for each experiment. Sections were then washed in PBS and incubated for 2 h at room temperature with species-appropriate Alexa Fluor–conjugated secondary antibodies. After final washes, sections were mounted, coverslipped with anti-fade mounting medium containing DAPI, and imaged on a Zeiss LSM700/780 confocal microscope or equivalent. For each biological replicate, 3–5 sections were analysed and 3–6 non-overlapping fields per section were quantified.

### RNAscope *in situ* hybridization

RNAscope *in situ* hybridization was performed on fixed mPFC sections using the RNAscope Multiplex Fluorescent Reagent Kit v2 (ACDBio, cat. no. 323270) according to the manufacturer’s instructions, with minor modifications. Mice were deeply anaesthetized with pentobarbital and transcardially perfused with ice-cold PBS followed by 4% paraformaldehyde in PBS. Brains were post-fixed overnight at 4 °C, cryoprotected in 30% sucrose, and sectioned coronally at 16 μm.

Sections were mounted on coated slides and processed using probes directed against *Grm5* (ACDBio, cat. no. 423631-C2), *Cnr1* (ACDBio, cat. no. 420721-C2), *Slc17a7* (*Vglut1*) (ACDBio, cat. no. 501101) and *Slc32a1* (*Vgat*) (ACDBio, cat. no. 319191-C3), as appropriate for each experiment. After protease treatment and probe hybridization, signal amplification and fluorescent detection were performed according to the kit protocol. Slides were counterstained with DAPI and coverslipped for imaging.

Images were acquired by confocal microscope (Zeiss LSM700/780) using identical acquisition settings within each experiment. Colocalization analyses were performed on anatomically matched mPFC fields. For each biological replicate, multiple sections and fields were analysed. Quantification was based on the fraction of marker-positive cells that co-expressed the target transcript, or conversely the fraction of target-positive cells that expressed the indicated marker, as specified in the figure legends.

### Image analysis

Images from immunofluorescence, RNAscope and histological verification experiments were analysed in Fiji/ImageJ or equivalent software. Identical acquisition settings were used within each experiment, and analysis was performed blind to condition where feasible. Linear adjustments of brightness and contrast were applied uniformly for display only and were not used for quantification.

## QUANTIFICATION AND STATISTICAL ANALYSIS

Statistical analyses were performed using MATLAB (R2021b or R2022a), GraphPad Prism 9, Origin 2018 and Microsoft Office 2019. Unless otherwise specified, data are presented as mean ± s.e.m. The statistical tests used for each experiment are indicated in the figure legends and described in the relevant sections of the Methods. Depending on experimental design, differences between groups were evaluated using two-sided Student’s *t* tests, one-way ANOVA or two-way repeated-measures ANOVA, with multiple-comparison correction where appropriate. Bonferroni adjustment was applied to ANOVA-based post hoc comparisons unless otherwise stated. Where standard parametric tests were not appropriate, custom permutation procedures were used as specified in the relevant analysis. Statistical significance was defined as *P* < 0.05. Exact *n* values are reported in the figure legends.

## Supplemental Figures and Legends

**Figure S1.**
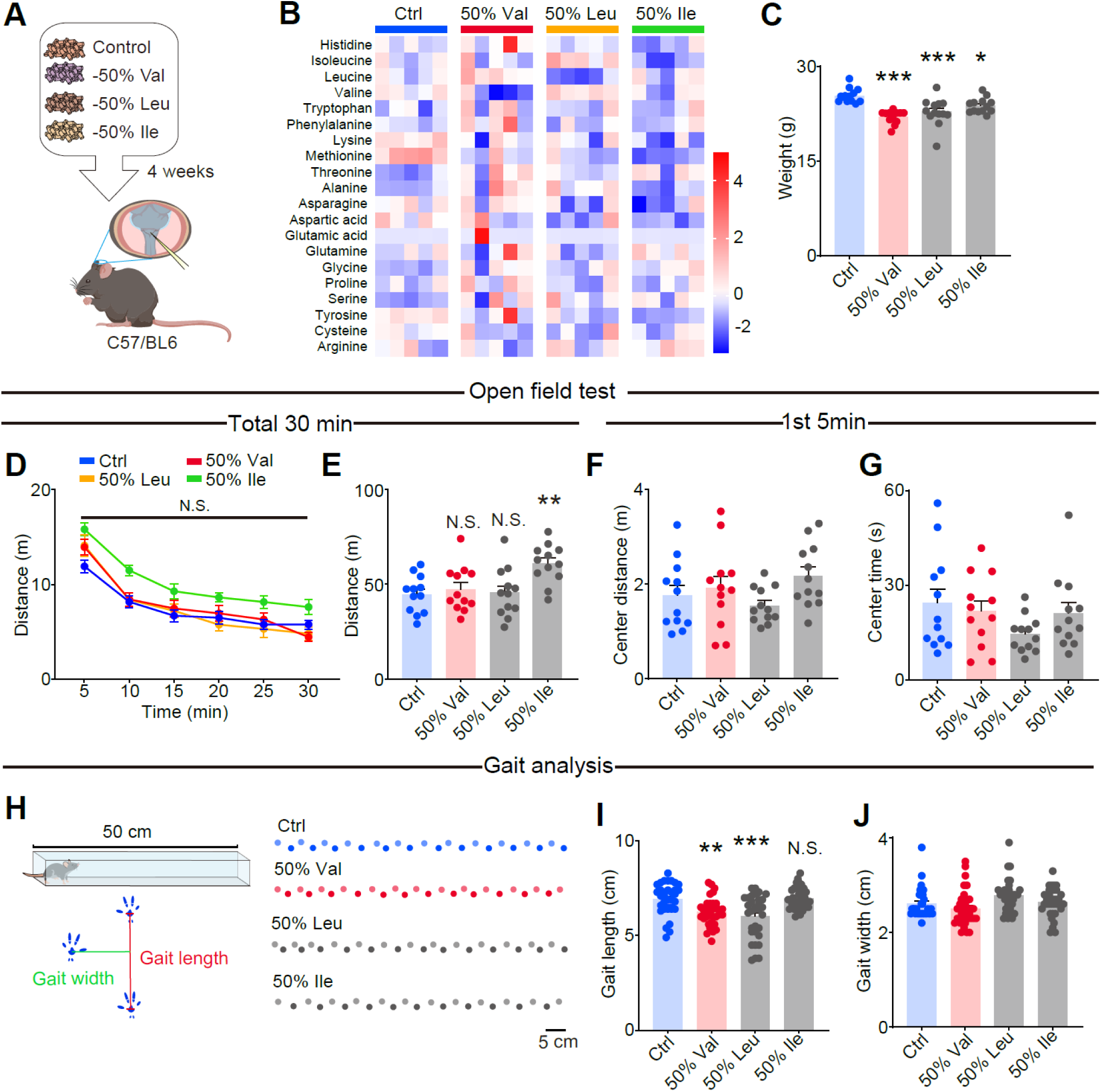
Systemic 50% restriction of individual BCAAs remodels CSF amino-acid profiles and produces growth and gait phenotypes, related to Figure 1. (A) Experimental design for dietary manipulation and CSF collection. Mice were maintained on control chow or diets in which valine, leucine or isoleucine was reduced by 50% for 4 weeks. (B) Heat map of CSF amino-acid metabolomics across diet groups (row-scaled), showing broad remodeling of the CSF amino-acid landscape under systemic BCAA restriction. (C) Body weight after dietary manipulation. *n* = 11–12 mice per group. (D–G) Open-field testing. (D) Distance travelled over time (30 min). (E) Total distance travelled over 30 min. (F and G) Center distance (F) and time spent in the center (G) during the first 5 min. *n* = 11–12 mice per group. (H) Gait analysis schematic (left) and representative footprints from each group (right). Scale bar, 5 cm. (I and J) Quantification of gait length (I) and gait width (J). *n* = 11–12 mice per group. Data are mean ± s.e.m.; dots represent individual mice. Statistics were analysed by one-way ANOVA with multiple comparisons, except for (D), which was analysed by two-way ANOVA as described in the Methods. \**P* < 0.05, \*\**P* < 0.01, \*\*\**P* < 0.001; N.S., not significant.

**Figure S2.**
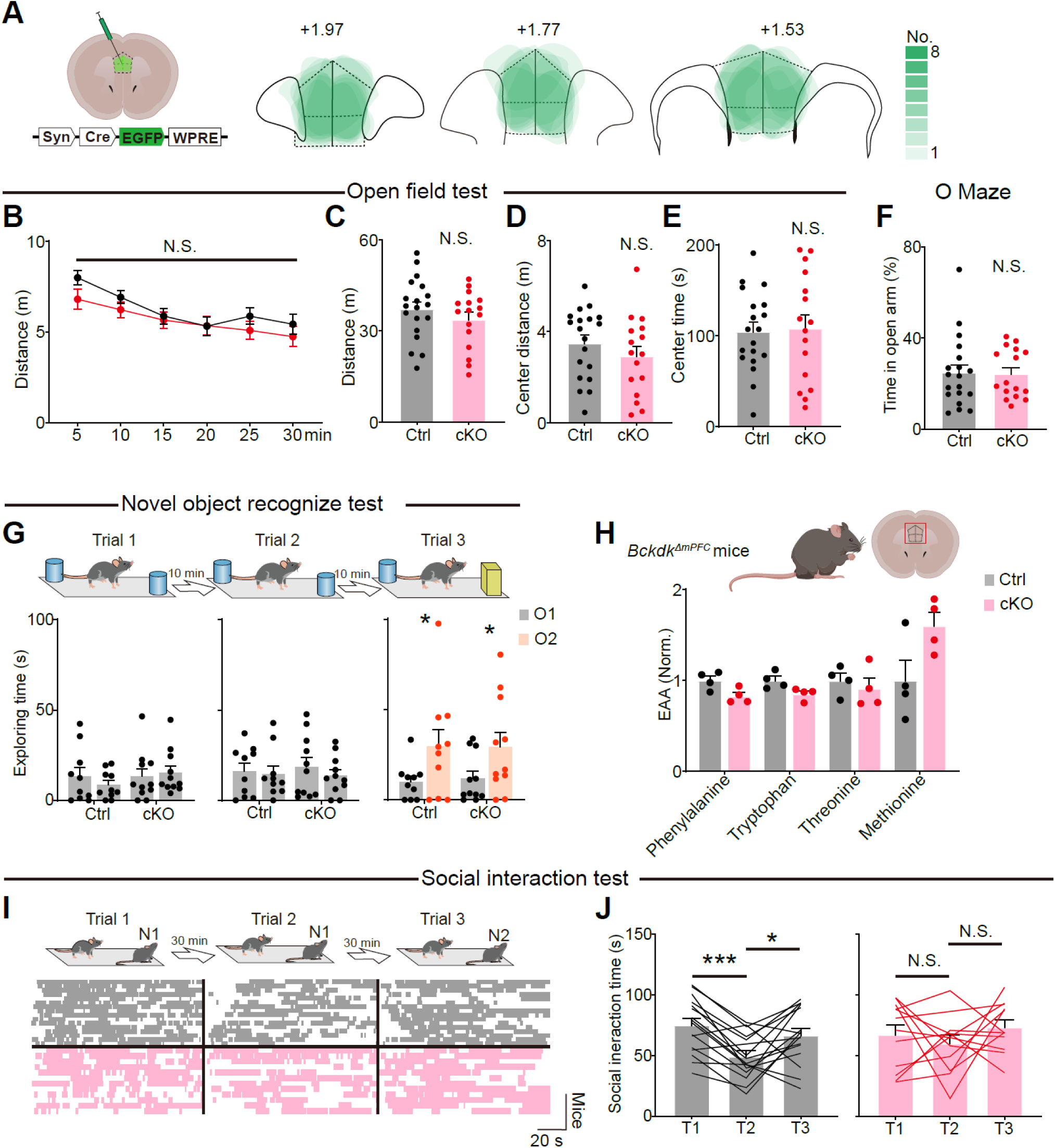
Generation and baseline phenotyping of *Bckdk*^ΔmPFC^ mice, related to Figure 1. (A) Bilateral mPFC delivery of AAV-Syn-Cre-EGFP-WPRE in *Bckdk^flox/flox^* mice. Heat maps summarize viral spread across animals at the indicated anterior–posterior coordinates relative to bregma; color scale denotes the number of mice contributing to each voxel. (B–E) Open-field testing. (B) Distance travelled over time (30 min). (C) Total distance travelled over 30 min. (D and E) Center distance (D) and time spent in the center (E) during the first 5 min. *n* = 16–19 mice per group, as shown. (F) Elevated O-maze: percentage time spent in the open arms. *n* = 16–19 mice per group, as shown. (G) Novel object recognition design (top) and quantification of object exploration across trials (bottom), indicating preserved object novelty preference. *n* = 11 mice per group. (H) Targeted profiling of selected non-BCAA essential amino acids in mPFC. Most are preserved, with changes as indicated. *n* = 4 biologically independent samples per group. (I) Direct social interaction assay design (top) and representative interaction rasters across trials (bottom): interaction with a novel conspecific (T1), re-exposure to the same conspecific (T2; habituation) and exposure to a new conspecific (T3; social novelty), separated by 30-min intervals. (J) Quantification of social interaction time across trials in control (left) and *Bckdk*^ΔmPFC^ (right) mice. *n* = 16 and 12 mice per group, as shown. Data are mean ± s.e.m.; dots represent individual mice (or biological samples in H). Statistical tests are described in the Methods. \**P* < 0.05, \*\**P* < 0.01, \*\*\**P* < 0.001; N.S., not significant.

**Figure S3.**
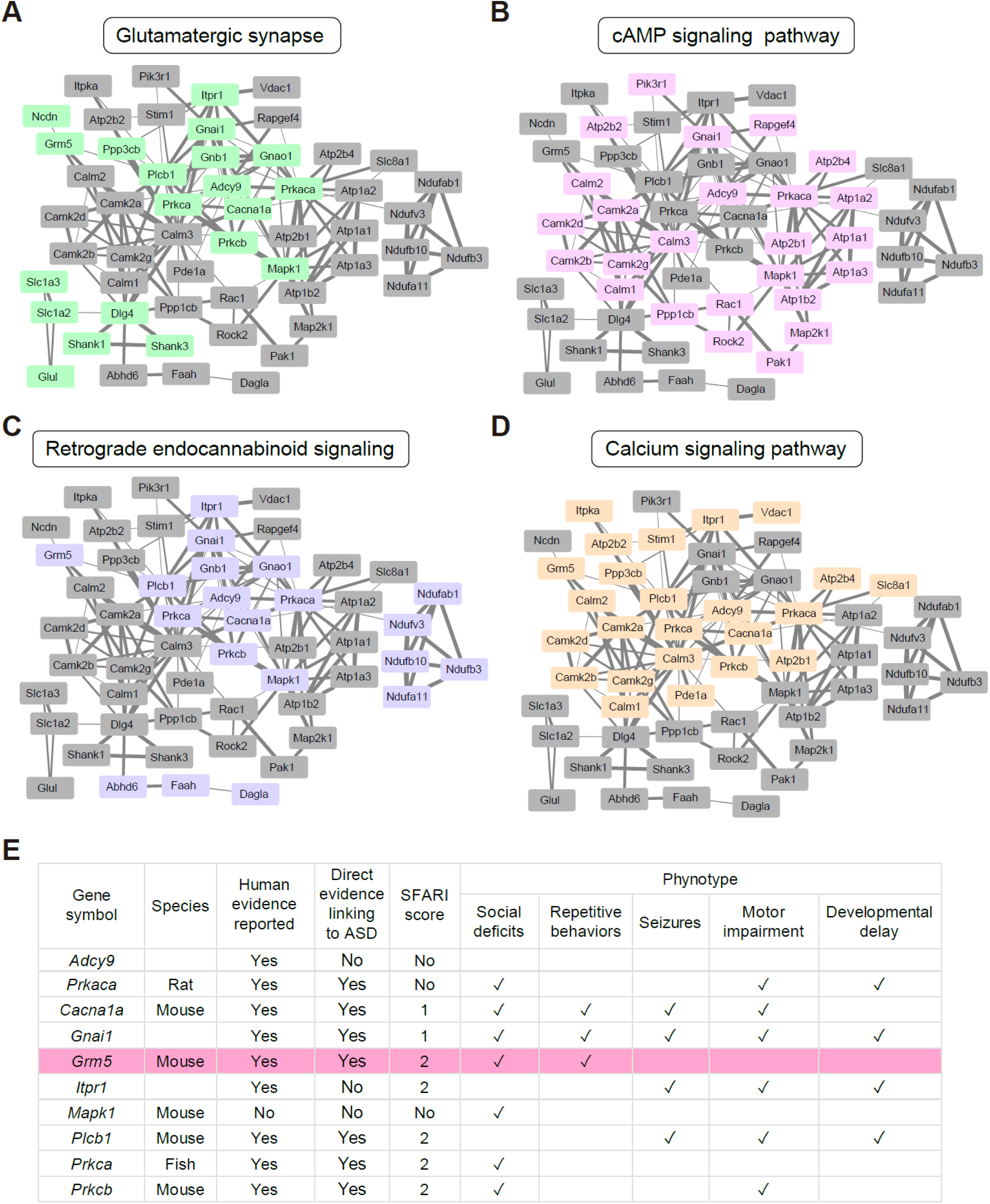
Pathway-resolved LiP-SMap candidates converge on synaptic signaling programs and highlight Grm5 as an ASD-relevant hub, related to Figure 2. (A–D) Protein interaction maps placing LiP-SMap-nominated candidates into four neurofunction-relevant pathways: glutamatergic synapse (A), cAMP signaling (B), retrograde endocannabinoid signaling (C) and calcium signaling (D). Highlighted nodes denote candidates with valine-dependent LiP-SMap signatures within each pathway module, emphasizing cross-pathway convergence. (E) Literature-based annotation of core multi-pathway nodes, summarizing reported human genetic or clinical links, direct experimental evidence relating each node to ASD-relevant phenotypes, SFARI classification (where applicable) and phenotype domains. *Grm5* is highlighted as a node with strong links to social phenotypes.

**Figure S4.**
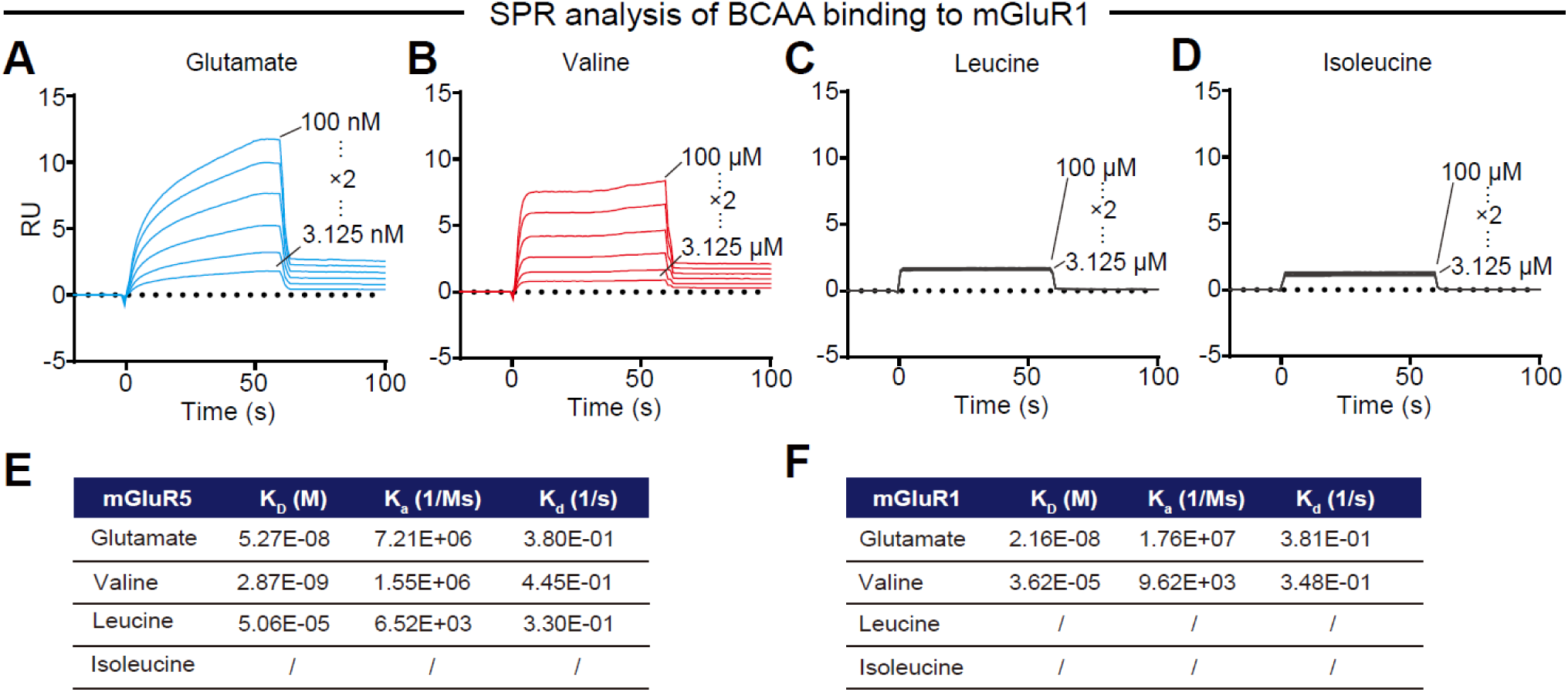
SPR defines receptor selectivity of BCAA binding, related to Figure 2. (A–D) SPR sensorgrams showing binding of glutamate (A), valine (B), leucine (C) and isoleucine (D) to mGluR1. Glutamate binds robustly in the nanomolar range, whereas valine binding is detected only at substantially higher concentrations; leucine and isoleucine show no detectable interaction under these conditions. (E and F) Kinetic parameters derived from SPR fitting for mGluR5 (E) and mGluR1 (F) (K_D_, K_a_, and K_d_), highlighting preferential high-affinity binding of valine to mGluR5 relative to mGluR1.

**Figure S5.**
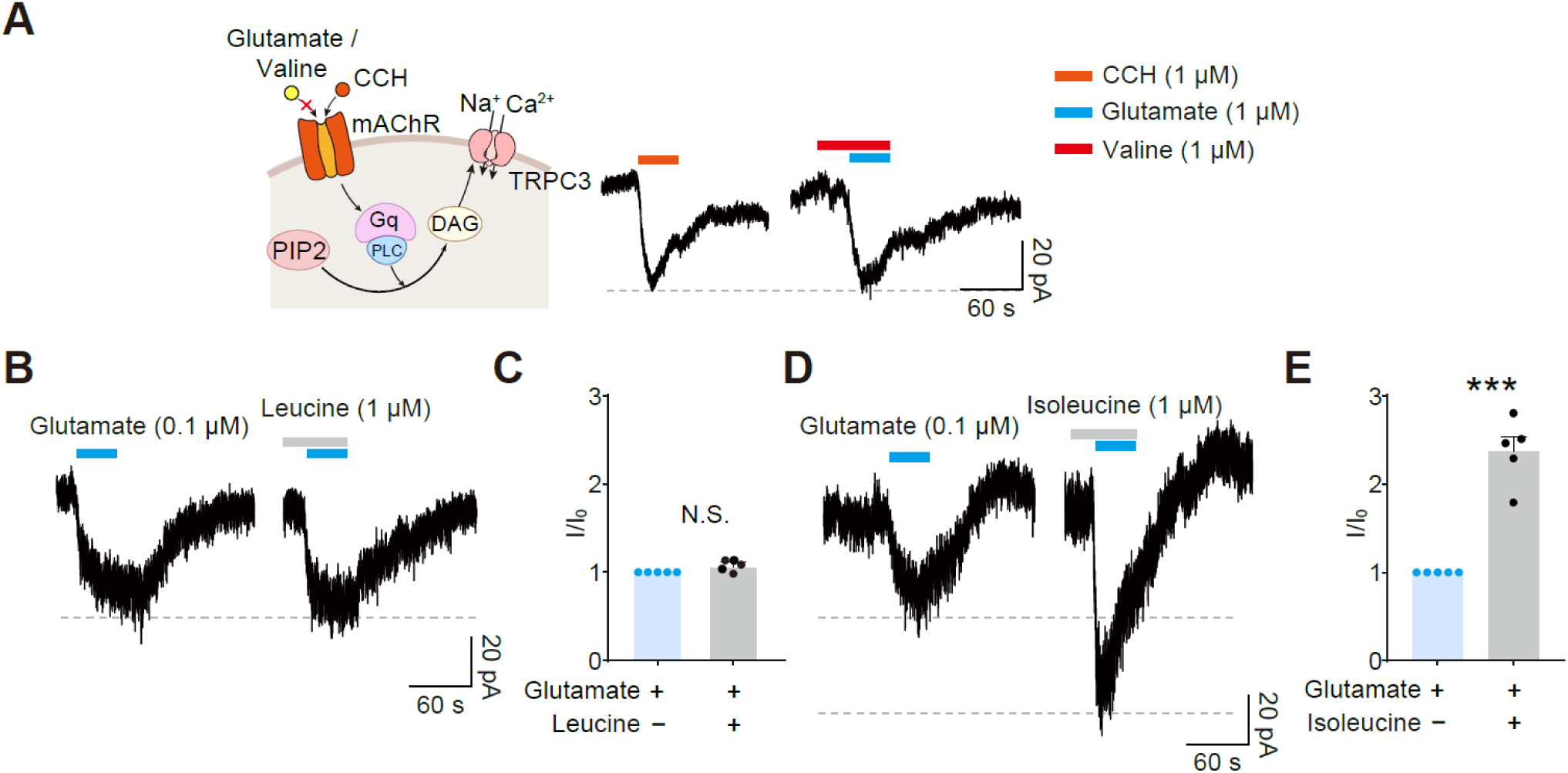
Additional electrophysiological analysis of valine-dependent potentiation, related to Figure 3. (A) Schematic of TRPC3 activation downstream of Gq-coupled receptors and representative recordings used to test whether valine acts on the downstream Gq–PLC–DAG–TRPC3 pathway independently of mGluR5. Activation of endogenous muscarinic receptors by carbachol (CCH) evokes TRPC3 current, whereas valine potentiation is assessed during mGluR5 stimulation. (B) Representative recordings showing that leucine pre-application does not potentiate glutamate-evoked TRPC3 currents. (C) Quantification of leucine-dependent potentiation of glutamate-evoked TRPC3 currents, expressed as a normalized current ratio (*I/I_0_*; see Methods). *n* = 5 cells per group. (D) Representative recordings showing that isoleucine pre-application produces a potentiation-like enhancement of glutamate-evoked TRPC3 currents. (E) Quantification of isoleucine-dependent potentiation of glutamate-evoked TRPC3 currents, expressed as a normalized current ratio (*I*/*I*_0_). *n* = 5 cells per group. Data are mean ± s.e.m. In (C) and (E), significance was assessed by two-sided unpaired Student’s *t* test. \**P* < 0.05, \*\**P* < 0.01, \*\*\**P* < 0.001; N.S., not significant.

**Figure S6.**
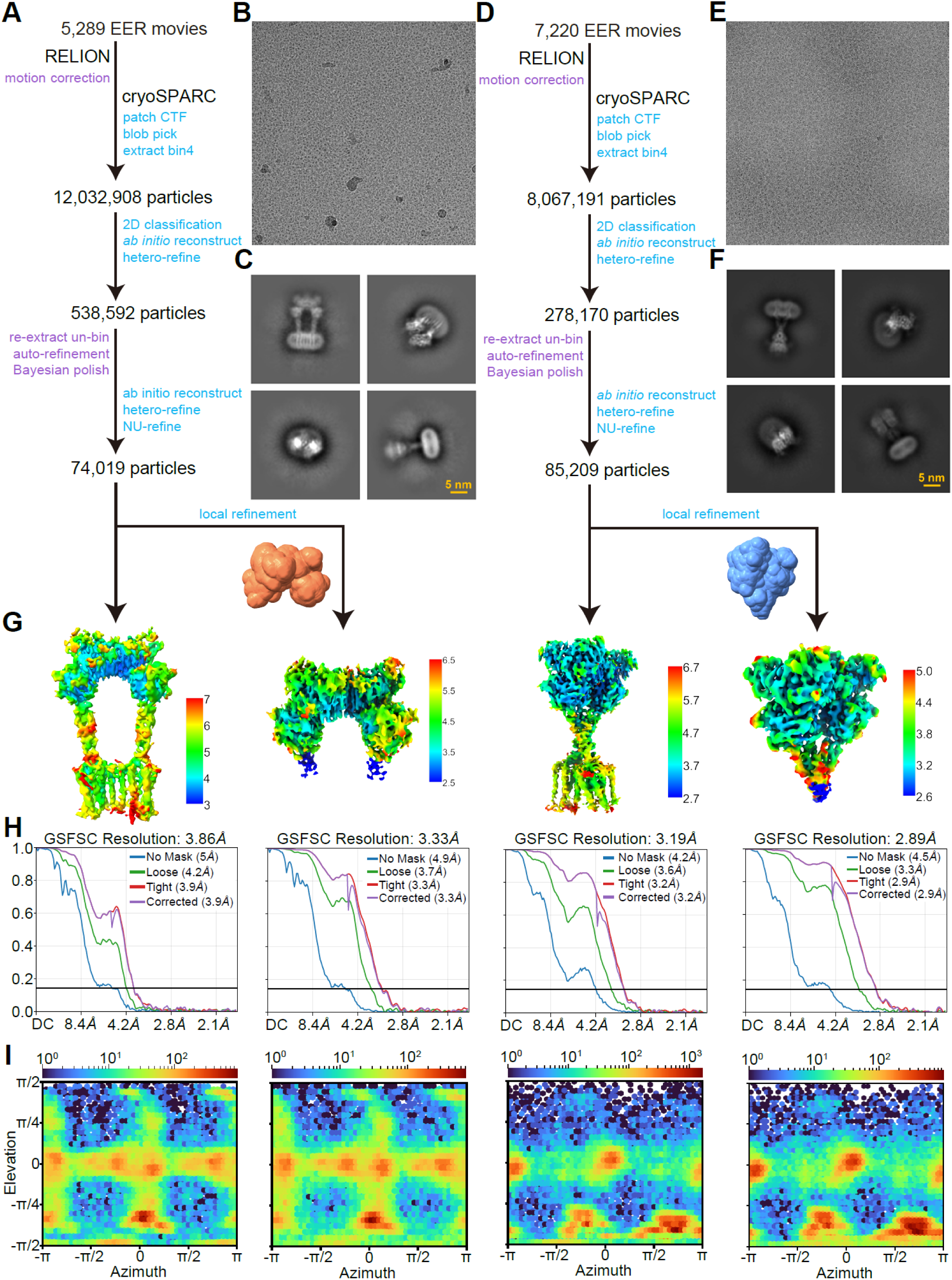
Cryo-EM processing and validation for apo-like and glutamate/valine-bound mGluR5 reconstructions, related to Figure 4. (A) Processing workflow for the apo-like mGluR5 dataset. A total of 5,289 EER movies were motion-corrected in RELION and processed in cryoSPARC (patch CTF estimation, particle picking, extraction, 2D classification, ab initio reconstruction, heterogeneous refinement and local refinement). Successive rounds of particle curation and refinement yielded a final global reconstruction from 74,019 particles. (B) Representative micrograph from the apo-like dataset. (C) Representative 2D class averages from the apo-like dataset. Scale bar, 5 nm. (D) Processing workflow for the glutamate/valine-bound mGluR5^GV^ dataset. A total of 7,220 EER movies were processed analogously, yielding a final reconstruction from 85,209 particles. (E) Representative micrograph from the mGluR5^GV^ dataset. (F) Representative 2D class averages from the mGluR5^GV^ dataset. Scale bar, 5 nm. (G) Final cryo-EM maps colored by local resolution for global reconstructions and VFT-focused local refinements. (H) Gold-standard FSC curves (0.143 criterion) for the global and locally refined maps: apo-like global map (3.86 Å), apo-like VFT local refinement (3.33 Å), mGluR5^GV^ global map (3.19 Å) and mGluR5^GV^ VFT local refinement (2.89 Å). (I) Angular distribution plots for the corresponding reconstructions.

**Figure S7.**
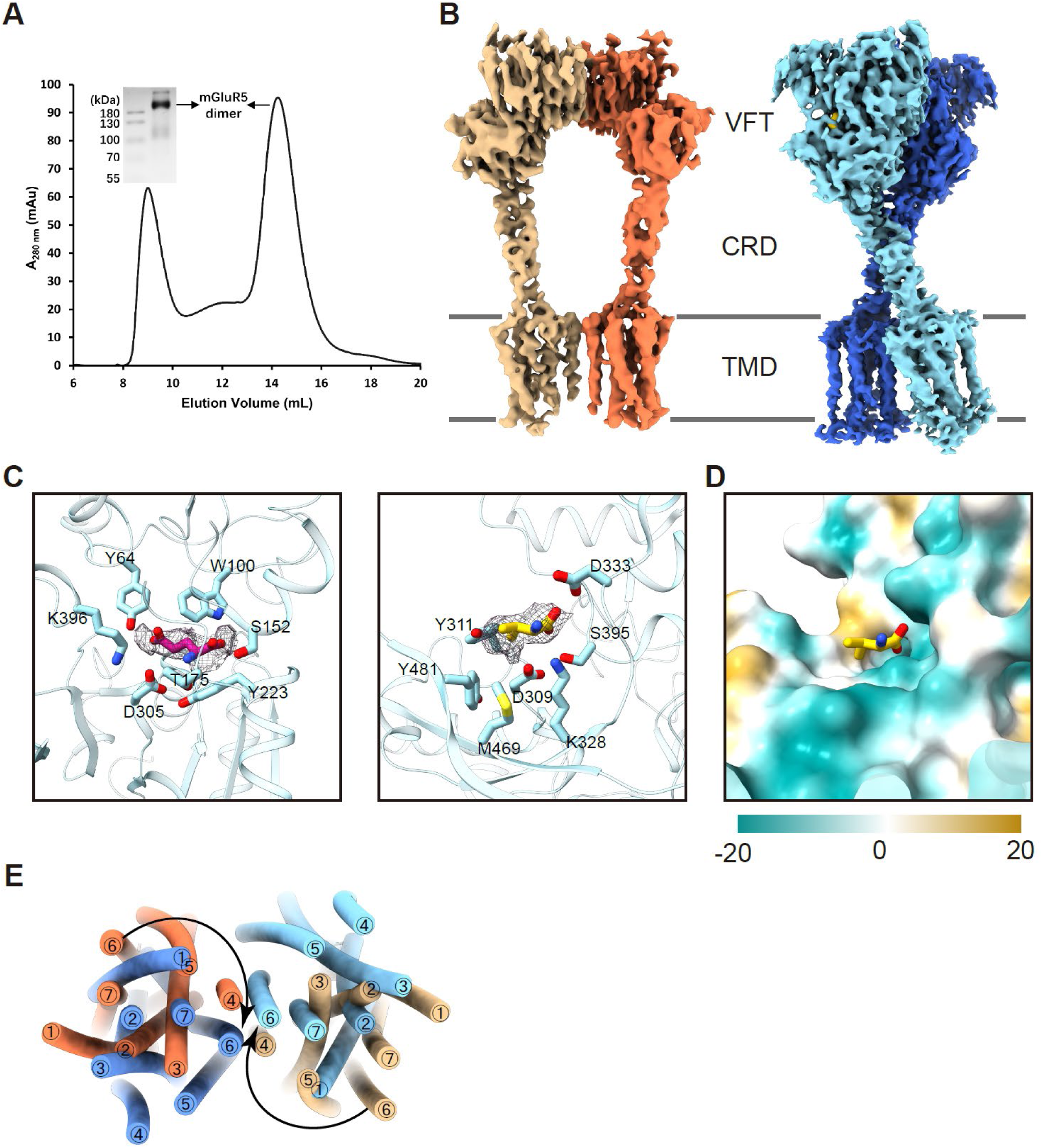
Purification, ligand density and activation-associated domain rearrangements in mGluR5 cryo-EM structures, related to Figure 4. (A) Representative size-exclusion chromatography profile of purified human mGluR5, with SDS–PAGE analysis shown in the inset. The major peak corresponds to a dimeric receptor preparation used for cryo-EM. (B) Cryo-EM density maps of apo-like mGluR5 (left) and glutamate/valine-bound mGluR5 (right), with each protomer colored separately. VFT, cysteine-rich domain (CRD) and transmembrane domain (TMD) are indicated. (C) Cryo-EM density for orthosteric glutamate (magenta, left) and allosteric valine (yellow, right) in the VFT domain. Ligand densities are shown as mesh at the same contour level in ChimeraX; coordinating residues are labelled. (D) Surface representation of the valine pocket colored by local physicochemical properties, highlighting a hydrophobic cavity accommodating the valine side chain together with flanking polar features that can engage the amino-acid backbone. (E) Schematic of TMD rearrangements comparing apo-like and glutamate/valine-bound mGluR5, illustrating closer approach of the two TMDs and formation of a TM6–TM6 interface in the ligand-bound state.

**Figure S8.**
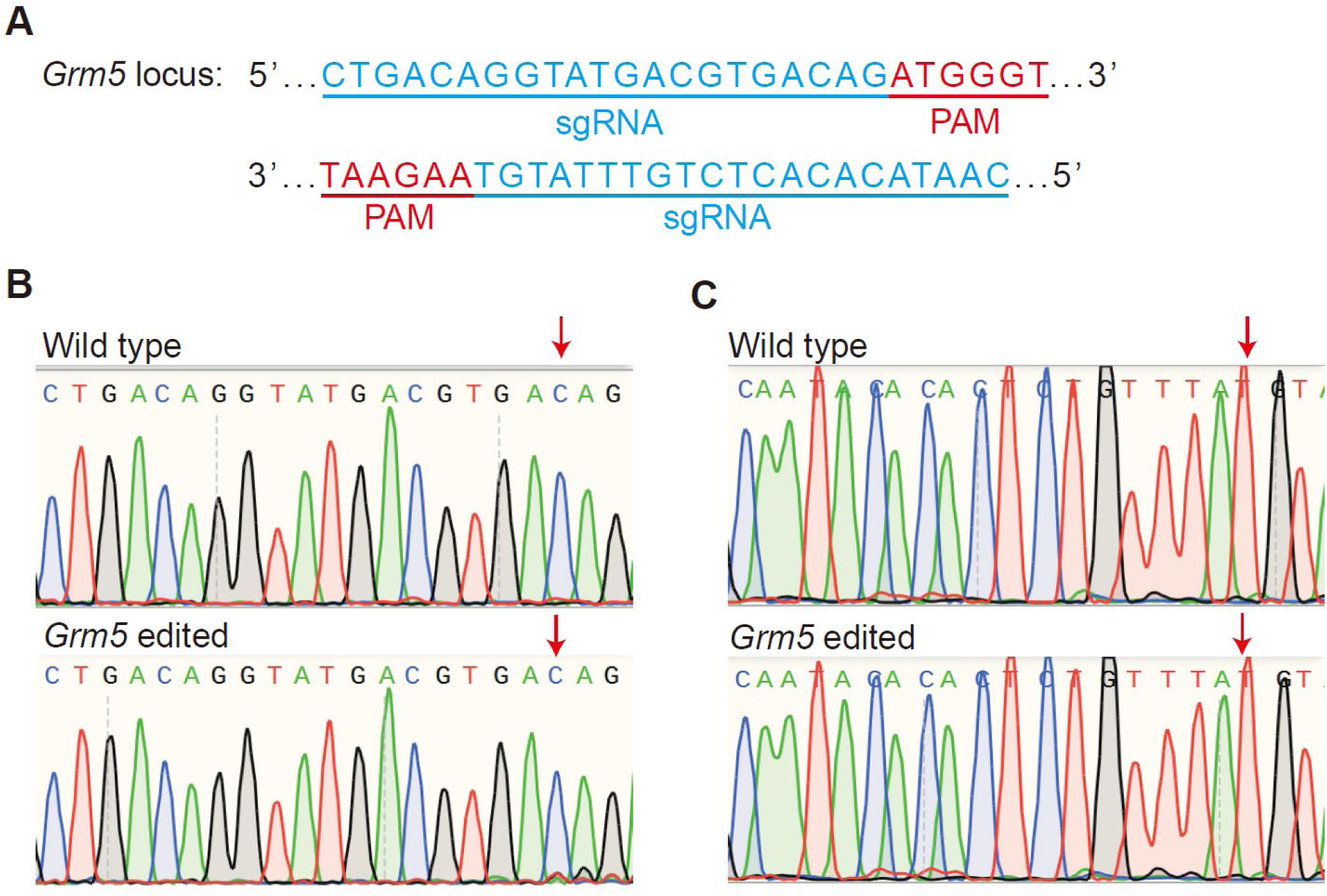
CRISPR–Cas9 targeting and validation of the *Grm5* locus, related to Figure 5. (A) Schematic of CRISPR–Cas9 targeting at the *Grm5* locus. The sgRNA target sequence is shown in blue and the adjacent PAM in red. Cas9 cleavage occurs approximately 3 bp upstream of the PAM, and a donor template was provided to enable homology-directed repair and introduction of the K327A substitution. (B and C) Representative Sanger sequencing chromatograms from wild-type and edited samples spanning the targeted genomic region, confirming successful editing at the *Grm5* locus.

**Figure S9.**
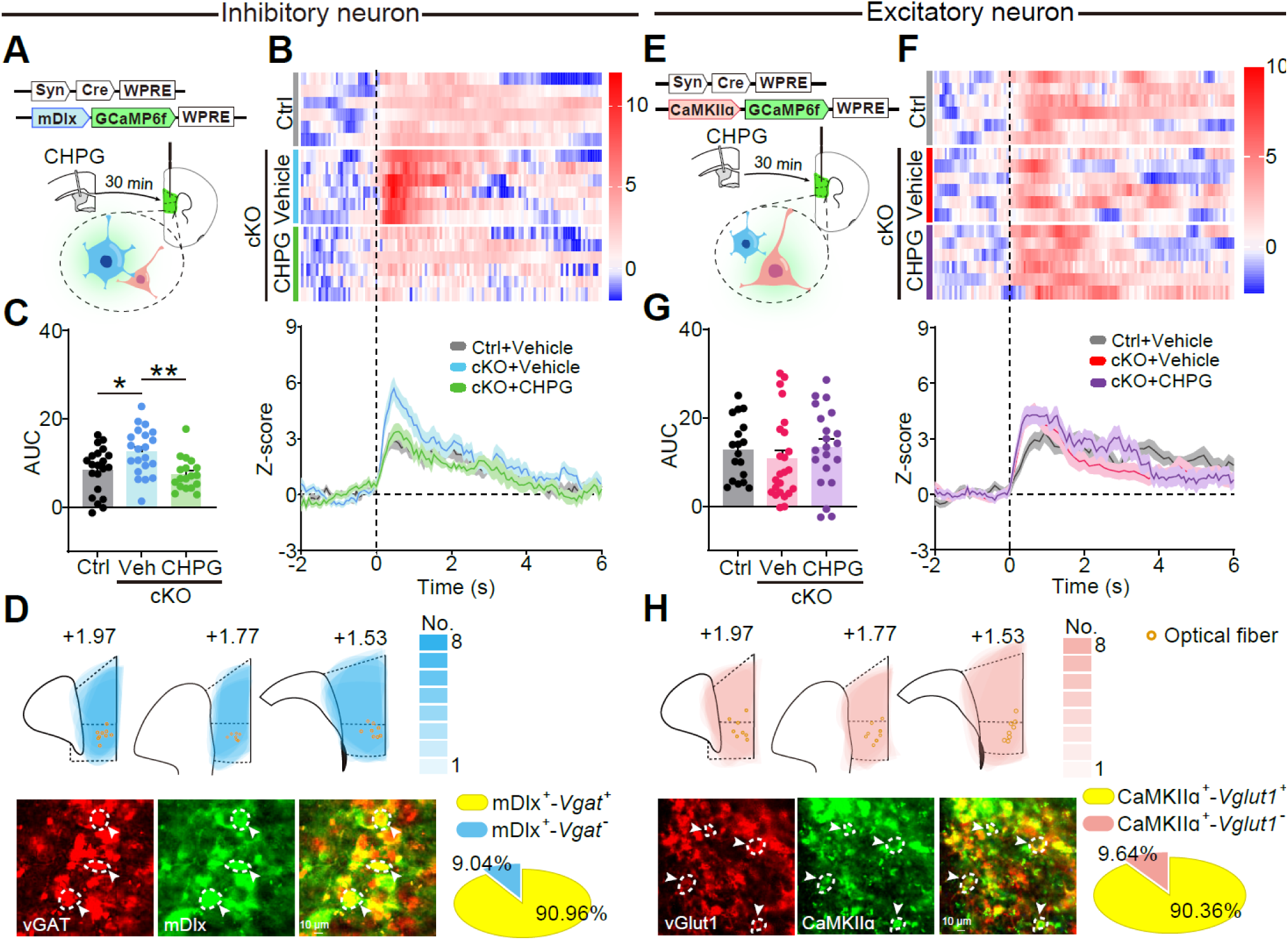
CHPG selectively normalizes social-evoked inhibitory-neuron activity in mPFC, related to Figure 6. (A) Experimental design for inhibitory neuron-biased fiber photometry in mPFC. Control and *Bckdk*^ΔmPFC^ (cKO) mice were injected in mPFC with an interneuron-enriched mDlx-GCaMP6f virus together with the viral strategy used to generate the local cKO, followed by optical fiber implantation; CHPG or vehicle was infused into mPFC 30 min before social interaction testing. (B) Social interaction-aligned z-scored *ΔF/F* signals from inhibitory-neuron recordings, shown as heat maps (left) and group-averaged traces (right) for Ctrl+Vehicle, cKO+Vehicle and cKO+CHPG. Time 0 indicates the onset of social investigation (dashed line). (C) Quantification of inhibitory-neuron Ca^2+^ responses as area under the curve (AUC) over the analysis window shown in (B). Each dot represents one social investigation bout; 18–21 bouts pooled from 6–7 mice per group. (D) Validation of inhibitory-neuron targeting. Top, mPFC coverage map across anterior–posterior coordinates with optical fiber locations indicated (orange). Bottom, representative co-labelling images for VGAT and mDlx and summary of targeting specificity (pie chart; percentages shown). Scale bar, 10 μm. *n* = 3 mice, 3 sections per mouse. (E) Experimental design for excitatory neuron-biased fiber photometry using CaMKIIα-GCaMP6f, in parallel with local CHPG or vehicle administration 30 min before testing. (F) Social interaction-aligned z-scored *ΔF/F* signals from excitatory-neuron recordings, shown as heat maps (left) and group-averaged traces (right). (G) Quantification of excitatory-neuron responses as AUC over the analysis window shown in (F). Each dot represents one social investigation bout; 18–24 bouts pooled from 6–8 mice per group. (H) Validation of excitatory-neuron targeting. Top, mPFC coverage map and optical fiber locations (orange). Bottom, representative co-labelling images for VGLUT1 and CaMKIIα and summary of targeting specificity (pie chart; percentages shown). Scale bar, 10 μm. *n* = 3 mice, 3 sections per mouse. Data are mean ± s.e.m. Multiple-group comparisons were analysed by one-way ANOVA with appropriate multiple-comparisons correction. \**P* < 0.05, \*\**P* < 0.01.

**Figure S10.**
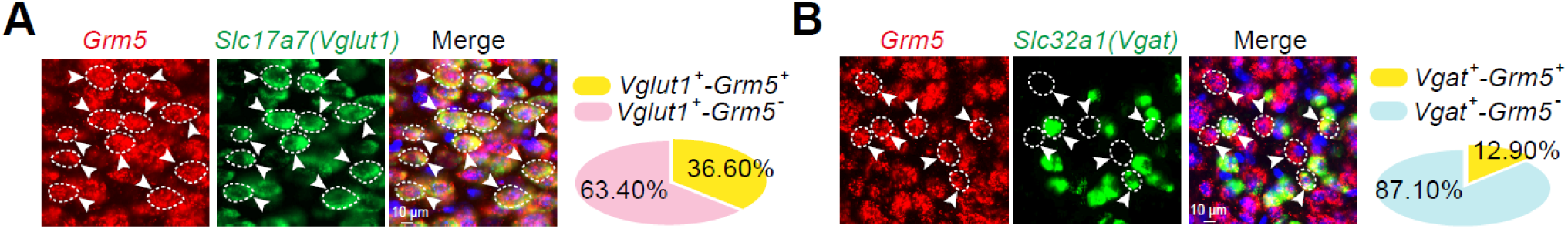
*Grm5* transcripts are more prevalent in excitatory than inhibitory neurons in mPFC, related to Figure 6. (A) Representative RNAscope images showing *Grm5* (red) co-localization with the excitatory marker *Slc17a7* (*Vglut1*, green) in mPFC. Dashed outlines indicate marker-positive cells and arrowheads indicate *Grm5* signal. Pie chart summarizes the fraction of *Vglut1*^+^ cells that are *Grm5*^+^. Scale bar, 10 μm. (B) Representative RNAscope images showing *Grm5* (red) co-localization with the inhibitory marker *Slc32a1* (*Vgat*, green) in mPFC, with corresponding quantification as in **a**. Pie chart summarizes the fraction of *Vgat*^+^ cells that are *Grm5*^+^. Scale bar, 10 μm. Quantification was performed across *n* = 3 mice, with 3 sections analysed per mouse.

**Supplemental Table 1.** Cryo-EM data collection and refinement statistics.

|  | **Apo mGluR5** | **mGluR5<sup>GV</sup>** |
| --- | --- | --- |
| **Data collection and processing** |  |  |
| Magnification | 130,000 | 130,000 |
| Voltage (kV) | 300 | 300 |
| Electron exposure (e<sup>-</sup>/Å<sup>2</sup>) | 50 | 50 |
| Defocus range (μm) | −1.2 to −2.0 | −1.2 to −2.0 |
| Pixel size (Å) | 0.932 | 0.932 |
| Symmetry | C1 | C1 |
| Initial particle (no.) | 12,032,908 | 8,067,191 |
| Final particle (no.) | 74,019 | 85,209 |
| Map resolution (Å) | 3.86 | 3.19 |
| **Refinement** |  |  |
| Initial model used (PDB code) | 8T6J | 8TAO |
| Map sharpening *B* factor (Å<sup>2</sup>) | 108.1 | 65.5 |
| Model composition |  |  |
| non-hydrogen atoms | 115520 | 11067 |
| Protein residues | 1528 | 1515 |
| Ligands |  | GGL:2;<br>VAL:1<br>NAG:2 |
| *B* factor (Å<sup>2</sup>) |  |  |
| Protein | 174.71 | 224.60 |
| Ligand |  | 90.18 |
| R.m.s. deviations |  |  |
| Bond lengths (Å) | 0.002 | 0.003 |
| Bond angles (°) | 0.551 | 0.672 |
| Validation |  |  |
| MolProbity score | 1.72 | 1.75 |
| Clash score | 9.67 | 9.30 |
| Poor rotamers (%) | 0.00 | 0.00 |
| Ramachandran plot |  |  |
| Favored (%) | 96.64 | 96.2 |
| Allowed (%) | 3.36 | 3.80 |
| Outliers (%) | 0.00 | 0.00 |
| Deposited model | 20YN | 9XFQ |
| Deposited map | EMD-67397 | EMD-66836 |

## Notes

### Competing Interest Statement

The authors have declared no competing interest.

